# RACK1 on and off the ribosome

**DOI:** 10.1101/582635

**Authors:** Alex G. Johnson, Christopher P. Lapointe, Jinfan Wang, Nicholas C. Corsepius, Junhong Choi, Gabriele Fuchs, Joseph D. Puglisi

**Author notes:** Contributed equally (Alex G. Johnson, Christopher P. Lapointe).

## Abstract

Receptor for activated C kinase 1 (RACK1) is a eukaryote-specific ribosomal protein implicated in diverse biological functions. To engineer ribosomes for specific fluorescent labeling, we selected RACK1 as an target given its location on the small ribosomal subunit and other properties. However, prior results suggested that RACK1 has roles both on and off the ribosome, and such an exchange might be related to its various cellular functions and hinder our ability to use RACK1 as a stable fluorescent tag for the ribosome. In addition, the kinetics of spontaneous exchange of RACK1 or any ribosomal protein from a mature ribosome *in vitro* remain unclear. To address these issues, we engineered fluorescently-labeled human ribosomes via RACK1, and applied bulk and single-molecule biochemical analyses to track RACK1 on and off the human ribosome. Our results demonstrate that, despite its cellular non-essentiality from yeast to humans, RACK1 readily re-associates with the ribosome, displays limited conformational dynamics, and remains stably bound to the ribosome for hours *in vitro*. This work sheds insight onto the biochemical basis of ribosomal protein exchange on and off a mature ribosome and provides tools for single-molecule analysis of human translation.

## INTRODUCTION

Protein synthesis by the ribosome is a heterogeneous and dynamic process across all kingdoms of life. As such, translation systems are particularly well-suited for single-molecule analysis, given its ability to resolve kinetics and parse out heterogenous pathways through real-time measurements. While nearly two decades of studies have clarified and expanded the mechanisms of bacterial translation, very few tools exist to study this process in human systems. A critical absence is an approach for high-efficiency labeling of human ribosomes with fluorescent dyes or other labels with minimal perturbation of ribosomal function or structure.

The eukaryotic ribosome contains ∼80 ribosomal proteins (RPs) and four strands of ribosomal RNA (rRNA) organized into large (60S) and small (40S) subunits (Melnikov et al. 2012). The ribosome is often considered a particle of stable composition, and it is thus surprising that nearly 20% of ribosomal proteins are non-essential in budding yeast (Steffen et al. 2012), implying that some RP-deficient ribosomes are capable of supporting cellular life. While much less is known about RP essentiality in mammals, preliminary work suggests that several RPs are non-essential within mammalian cell lines (Blomen et al. 2015) (Supp. Fig. 1a). This realization was our gateway into the fluorescent labeling of human ribosomes, wherein we leveraged the cellular non-essentiality of eS25 to genetically engineer a cell line bearing ribosomes labeled at that protein (Fuchs et al. 2015). However, this method has several limitations: in particular it relies on a large protein tag (SNAP-tag) that more than doubles the size of eS25, which may obscure analyses of conformational dynamics and reduce ribosome labeling efficiency through misfolding or proteolysis of the fusion protein. We were further concerned that such a system might artificially perturb or fail to monitor some translation events given the location of eS25 near the ribosomal E site and that sub-populations of cellular ribosomes are sub-stoichiometric for eS25 (Shi et al. 2017; Van De Waterbeemd et al. 2018).

To address these issues, our focus turned to another eukaryote-specific ribosomal protein: Receptor for activated C kinase 1 (RACK1). Given its prominent position at the surface-exposed region of the 40S ribosomal subunit (Fig. 1a), we predicted that labeling at RACK1 would be less likely to perturb translation events. Nonetheless, RACK1 has been implicated in diverse biological roles (Gibson 2012), and we were concerned that its labeling might inadvertently alter ribosome function. For one, RACK1 is a WD40-domain protein, a domain seen broadly in cellular signaling proteins and often implicated in a “scaffolding” role (Stirnimann et al. 2010). While RACK1 is non-essential for cellular life from yeast to humans (Jha et al. 2017) (although a full knockout is embryonically lethal in mice (Volta et al. 2013)), its loss or depletion leads to altered cellular signaling (Gibson 2012; Nielsen et al. 2017), translational rewiring (Thompson et al. 2016; Gallo et al. 2018), misregulated ribosome-associated quality control (Sundaramoorthy et al. 2017; Juszkiewicz and Hegde 2017; Sitron et al. 2017; Matsuo et al. 2017), and protection against multiple viruses (Majzoub et al. 2014; Jha et al. 2017; Kim et al. 2018; Hafirassou et al. 2017). These functional aspects of RACK1 suggest a need for a minimal perturbation of RACK1 for labeling and subsequent analyses to demonstrate that cellular function is not perturbed.

**Figure 1.**
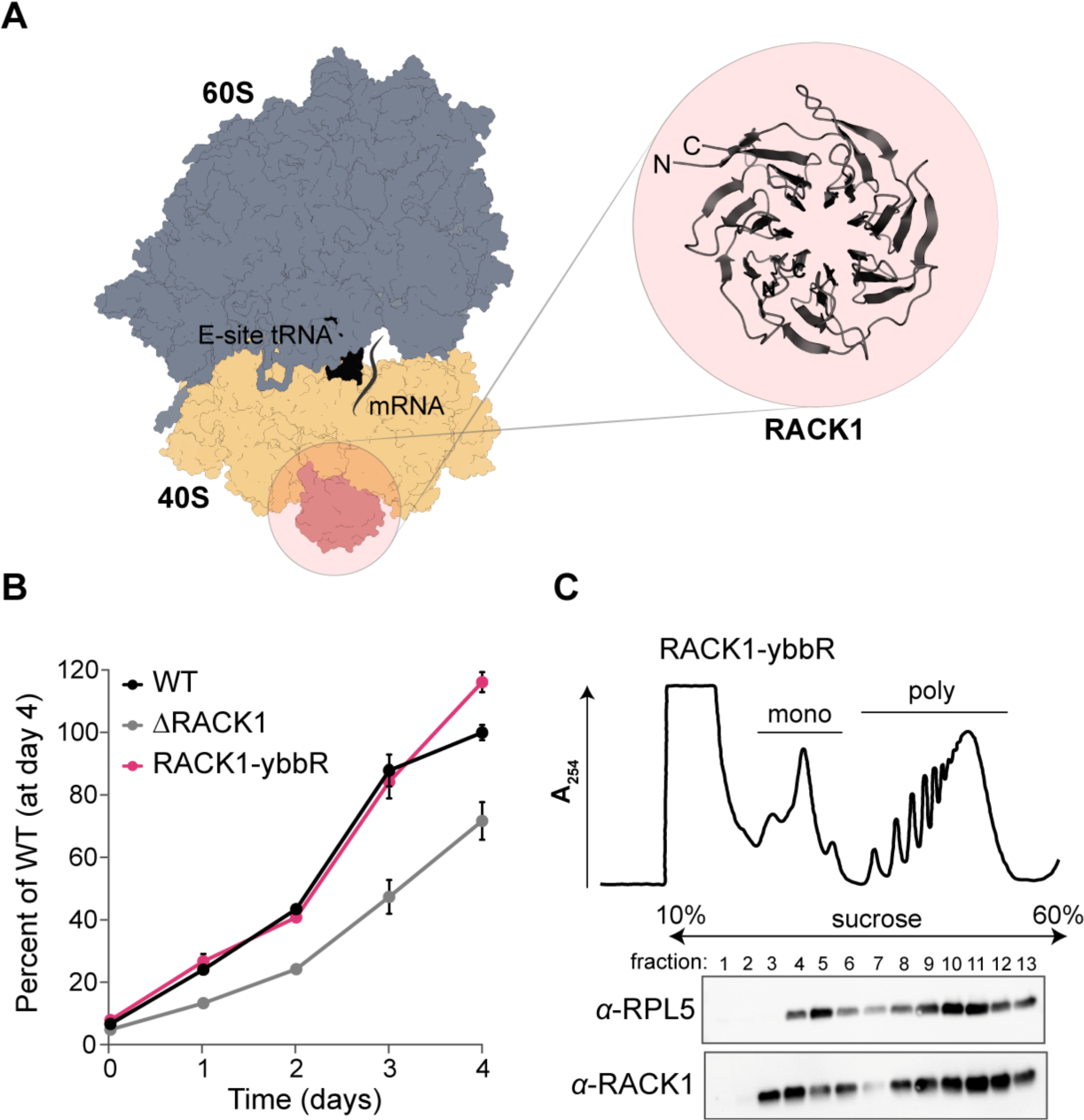
RACK1-ybbR incorporates into translating ribosomes in human cells. **A.** Structure of the human ribosome (PDB 6ek0). The 40S (tan) and 60S (blue) ribosomal subunits, RACK1 (pink), mRNA (grey line) and an E-site tRNA (black) are indicated. The WD40 repeat structure of RACK1 is illustrated in the inset with a ribbon depiction (also from PDB 6ek0). There are one and three unresolved amino acids on the N- and C-termini of RACK1, respectively. **B.** Plot of the proliferation of the indicated cell lines as measured by an MTT cell proliferation assay relative to the wild-type cells at day 4. Error bars represent the 95% CI (*n* = 10). **C.** UV absorbance trace (at 254 nm) of polysome profiling and associated western-blot analysis of the RACK1-ybbR cell line. “Mono” and “poly” refer to peaks that correspond to monosomes and polysomes, respectively.

Another confounding factor for labeling ribosomes at RACK1 originates from cellular studies that attempt to distinguish the ribosomal and non-ribosomal functions of the protein (Gibson 2012; Nielsen et al. 2017). Confusion here is warranted—in stationary phase yeast, the RACK1 homolog Asc1p is detected free in the cytosol (Baum et al. 2004), and some of its properties listed above seem to suggest an extra-ribosomal role (Warner and McIntosh 2009). The potential for RACK1 exchange from the ribosome is also suggested by it being one of the final proteins to incorporate during ribosome biogenesis (Larburu et al. 2016), yet it is also a stoichiometric component of most eukaryotic ribosomes studied to date (Sengupta et al. 2004; Rabl et al. 2011; Shi et al. 2017; Van De Waterbeemd et al. 2018). However, a notable exception is in *Plasmodium falciparum* (the malaria parasite) where RACK1 is expressed and required for intracellular survival (Blomqvist et al. 2017), but paradoxically not present in any ribosome structure (Wong et al. 2014; Sun et al. 2015). Thus, RACK1 embodies suggestive qualities of an exchangeable ribosomal protein, and these properties could be intimately linked with its function.

Here our first goal was to establish a stable and stochiometric human ribosome labeling site, but given the uncertain stability of RACK1 on the ribosome and the unique opportunity it provided, we sought to harness RACK1 to study the biochemical basis for ribosomal protein exchange. While a few studies have demonstrated the possibility for RP incorporation into ribosomes lacking an RP (Lieberman et al. 2000; Cornish et al. 2008; Kossinova et al. 2008), the kinetics of this process have not been investigated in the context of mature ribosomes *in vitro*. After labeling human ribosomes at RACK1 and thereby further facilitating biophysical analyses of translation, we devised a system to track RACK1 flux on and off the ribosome to address this knowledge gap. Collectively, our results demonstrate the benefit of compact and targeted labeling schemes for the human ribosome, which is generalizable to other RPs. By following the dynamics of RACK1 association to and dissociation from ribosomes with single-molecule spectroscopy, we further establish an *in vitro* benchmark for the biochemical stability of an integral ribosomal protein.

## RESULTS

RACK1 is organized into a symmetric and well-folded structure when free in solution (Fig. 1a) as revealed by crystallographic studies on proteins expressed in and purified from *E. coli* (Bjørndal et al. 2003; Nery et al. 2006; Coyle et al. 2009; Carrillo 2012). Whereas most other RPs are small and very basic (pI ≈ 10-11), RACK1 has a relatively large molecular weight and near neutral pI (Supp. Fig. 1b,c). In our hands, the non-essential ribosomal proteins eS25 and eL22 may be expressed in *E. coli* when fused to maltose-binding protein (MBP), but readily precipitate at physiological salt concentrations. Thus, based on its biochemical tractability and cellular non-essentiality, we developed a strategy to fluorescently label RACK1 on and off the ribosome as a strategy to label intact ribosomes and to monitor its ribosome association as a free protein. To ensure a compact labeling strategy, we utilized site-specific labeling with the 11 amino acid ybbR-tag (Yin et al. 2006) (Fig. 1a).

We first tested whether ybbR-tagged RACK1 is functional on ribosomes in cell culture. Using lentiviral transduction, we expressed C-terminally ybbR-tagged RACK1 in a previously-generated knockout of RACK1 (ΔRACK1) in the human HAP1 cell line (Jha et al. 2017), and examined wild-type, ΔRACK1, and RACK1-ybbR cell lines using cell proliferation and polysome profiling assays. Whereas the ΔRACK1 cell line grows slower relative to the wild-type cell line (Fig. 1b and Supp. Fig. 2a), expression of RACK1-ybbR in the ΔRACK1 cell line recovered cell proliferation to wild-type levels. To probe its functionality, we examined the three cell lines using polysome profiling and western blot analyses. RACK1 and RACK1-ybbR were almost completely associated with ribosomes, with minimal free protein detected, and the proteins were present in both monosome and polysome fractions (Fig. 1c and Supp. Fig. 2b). In the ΔRACK1 cell line, we observed an increase in the monosome peak relative to the polysome peaks, possibly indicating alterations in the pool of active ribosomes (Supp. Fig. 2b). Thus, RACK1-ybbR rescues the ΔRACK1 growth defect and is incorporated into actively translating ribosomes.

### Fluorescent labeling of intact 40S ribosomal subunits via RACK1-ybbR

RACK1 is a prominent feature on the surface-exposed region of the 40S subunit, where it contacts ribosomal proteins uS3, uS9, and eS17, and helices 39 and 40 of 18S rRNA, and may facilitate interactions with numerous signaling proteins (Nielsen et al. 2017). It is proximal to the mRNA entry and exit channels (Fig. 2a), and adjacent to the region bound by the HCV IRES, which directly binds the 40S subunit with high-affinity (Johnson et al. 2017). The N- and C-termini of RACK1 are in close proximity and surface accessible, so the C-terminal ybbR-tag on RACK1 should be accessible to SFP synthase (Yin et al. 2006) on the mature 40S ribosomal subunit (Fig. 2b,c). To test this, we purified ribosomes from RACK1-ybbR expressing cells, identified optimal conditions for labeling (Supp. Fig. 3a,b), and routinely achieved labeling efficiencies >50% on purified ribosomal subunits.

**Figure 2.**
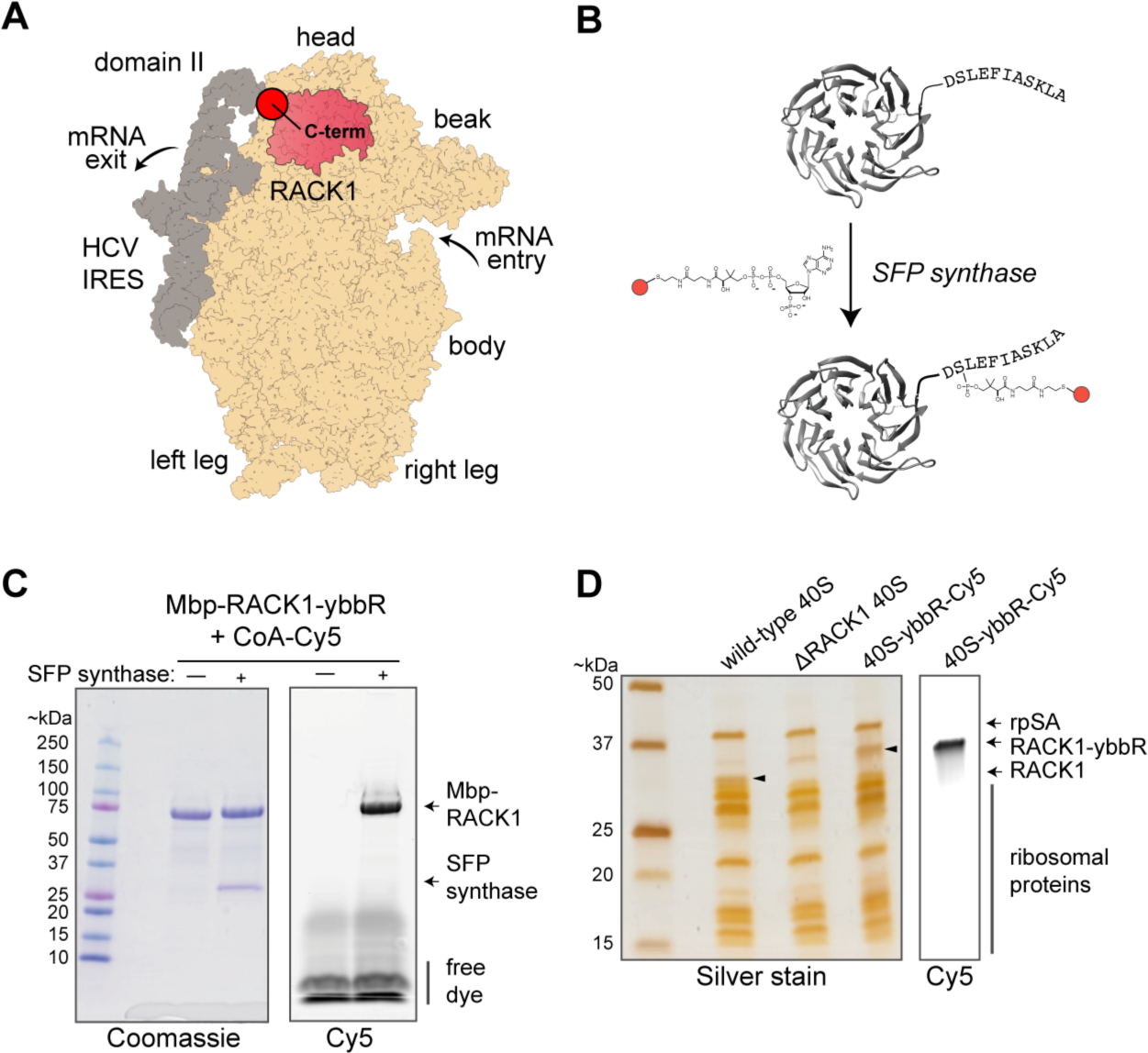
Strategy for labeling ribosome-bound and recombinant RACK1-ybbR with fluorophores using SFP synthase. **A.** Model structure of the human 40S ribosomal subunit (tan) bound to the HCV IRES (grey), with RACK1 highlighted in pink (PDB 5a2q). Key locations on the ribosome and IRES are indicated, with this view focused on the solvent-exposed surface of the 40S subunit. **B.** Model of human RACK1 structure with the C-terminal ybbR peptide tag indicated. RACK1-ybbR was site-specifically labeled by incubation with SFP synthase and CoA-Cy3 or CoA-Cy5 conjugated dye substrates. **C.** Image of a representative gel that was first scanned for Cy5 fluorescence (right) and subsequently stained with Coomassie blue (left) following SDS-PAGE analysis of RACK1-ybbR post-labeling with and without SFP synthase. **D.** Image of a representative gel that was first scanned for Cy5 fluorescence (right) and subsequently silver stained (left) following SDS-PAGE analysis of the indicated 40S ribosomal subunits, with the RACK1 bands indicated by arrows on both images.

To determine whether RACK1-ybbR containing 40S ribosomal subunits were biochemically intact, we characterized them alongside wild-type and ΔRACK1 subunits using denaturing and native gel assays. By SDS-PAGE, we observed a single fluorescent band in the 40S-RACK1-ybbR sample that matched the migration pattern of recombinant RACK1-ybbR (Fig. 2d and Supp. Fig. 3c). This RACK1 band with an increased molecular weight was detected in 40S ribosomal subunits at a level similar to endogenous RACK1 in wild-type 40S subunits by immunoblotting (Supp. Fig. 3d). As expected, no RACK1 was detected in the ΔRACK1 40S subunits. Several other ribosomal proteins also exhibited similar levels across the three 40S samples; however, we observed a modest increase in a higher molecular weight band for uS3 in ΔRACK1 relative to wild-type 40S subunits, which was returned to near wild-type levels in 40S-RACK1-ybbR subunits (Supp. Fig. 3d). The C-terminus of uS3 directly contacts RACK1, and this band may represent a post-translational modification to uS3 that was increased in the absence of RACK1. Finally, using native-gel electrophoresis (Johnson et al. 2018), we found that 40S-RACK1-ybbR subunits bound the HCV IRES with similar efficiency as to WT 40S subunits, and they were competent to form Mg^2+^-driven (Lancaster et al. 2006; Yamamoto et al. 2014) 80S complexes on the IRES (Supp. Fig. 3e). In summary, 40S-RACK1-ybbR ribosomes are nearly identical in composition to wild-type subunits and functional in preliminary biochemical assays.

### A reliable smFRET signal for the presence of RACK1 within 40S subunits

Given the high-affinity between the 40S subunit and HCV IRES and its utility for specific immobilization of human ribosomes (Fuchs et al. 2015; Johnson et al. 2018), we next examined the suitability of 40S-RACK1-ybbR ribosomal subunits for single-molecule analyses in the context of the HCV IRES. Domain II of the IRES is predicted to be within smFRET distance with the C-terminus of RACK1, despite there being no direct RNA-RACK1 interaction (Fig. 3a). We therefore used segmental labeling to attach fluorophores directly to the HCV IRES to generate two versions of the IRES with Cy5 dyes at either the base (C44) or elbow (U56) of domain II (Supp. Fig. 4a). These positions are predicted to be within the dynamic range of FRET (about 70 and 90 Å away from the C-terminus of RACK1 for C44 and U56, respectively (Quade et al. 2015; Yamamoto et al. 2014); Fig. 3a), and allow an approximate triangulation of conformational dynamics within the 40S-IRES complex.

**Figure 3.**
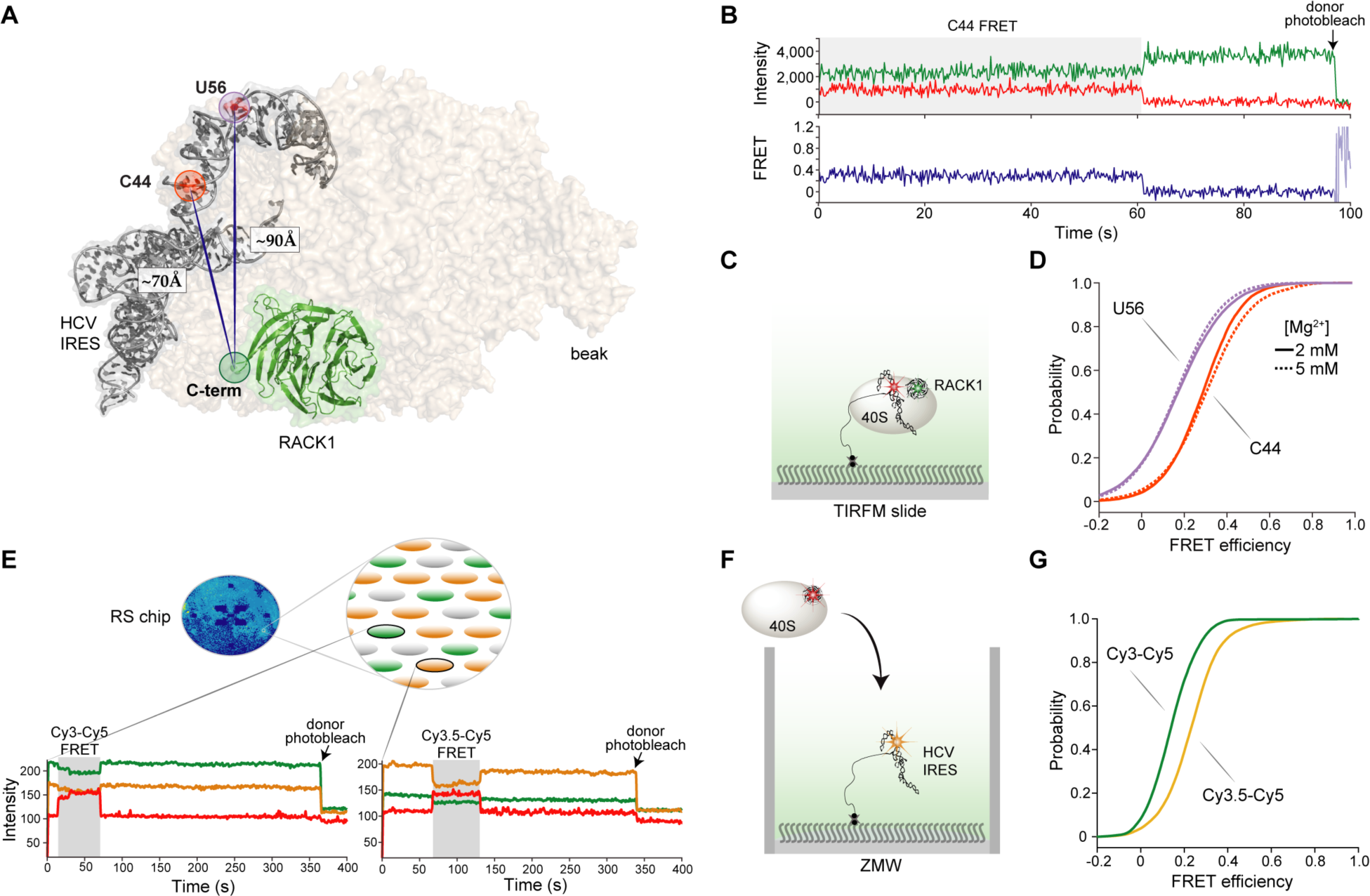
RACK1-ybbR is proximal to domain II of the HCV IRES and provides a signal for 40S-IRES association and conformational dynamics on two single-molecule fluorescence platforms. **A.** Structural model of the human 40S ribosomal subunit (tan) bound to the HCV IRES (grey), with RACK1 highlighted in green (PDB 5a2q). The C-terminus of RACK1, and the locations of C44 and U56 in the HCV IRES are highlighted, with the measured distance between them indicated. **B.** Representative single-molecule fluorescence trace and FRET conversions from TIRFM experiments, with FRET between RACK1-ybbR-Cy3 and IRES(C44-Cy5) highlighted in grey. smFRET was signified by anti-correlated changes in fluorescence intensity of the fluorophores, and single 40S-IRES complexes were identified via a single-step photobleach in the Cy3 donor signal, indicated by the arrow. **C.** Schematic of surface immobilized 40S-RACK1-ybbR-Cy3:Cy5-HCV-IRES complexes on a quartz slide used for TIRFM. **D.** Cumulative distribution plot of observed FRET intensities for 40S-RACK1-ybbR-Cy3 with IRES(Cy5-U56, purple) or IRES(Cy5-C44, orange) in the presence of 2 mM (solid lines) or 5 mM Mg^2+^ (dashed lines). The number of traces analyzed (*n*) for panels d are listed in Supplementary Figure 4i. **E.** Schematic of 40S-RACK1-ybbR-Cy5 delivery to surface-immobilized HCV IRES labeled at C44 with Cy3 or Cy3.5 in zero-mode waveguides (ZMWs). The blue oval is the full view of an RS chip that has both IRES(Cy3-C44) and IRES(Cy3.5-C44) immobilized in individual ZMWs on the chip. Below the schematic are example single-molecule fluorescence traces that depict Cy3-Cy5 (left) or Cy3.5-Cy5 (right) upon delivery of 40S-RACK1-ybbR-Cy5 to dual-immobilized IRES(Cy3-C44) and IRES(Cy3.5-C44). The respective FRET events are highlighted in gray, and photobleaching of the donors are indicated by the arrows. **F.** Schematic of an individual ZMW showing delivery of 40S-RACK1-ybbR-Cy5 to a Cy3.5 labeled HCV IRES. **G.** Cumulative distribution plots of observed FRET efficiency from 40S-RACK1-ybbR-Cy5 FRET with either IRES(Cy3-C44) (green) or IRES(Cy3.5-C44) (yellow) after delivery to dual-immobilized IRESs. Both data sets were fit with single Gaussian distributions. The number of traces analyzed (*n*) for panel g are listed in Supplementary Figure 5d.

To detect smFRET between RACK1 and the HCV IRES, we immobilized complexes on a Neutravidin-coated quartz slide and imaged them using total internal reflection fluorescence microscopy (TIRFM) and laser excitation of Cy3-ybbR-RACK1 on the 40S subunit (Fig. 3b,c and Supp. Fig. 4b). We observed smFRET to Cy5-IRES(C44) or Cy5-IRES(U56) in 40S-IRES complexes (Fig. 3b). The resulting smFRET signals from both dye pairs were characterized at near- and above-physiological Mg^2+^ concentrations (2 and 5 mM, respectively) (Fig. 3d and Supp. Fig. 4c-g) and in a Mg^2+^-driven 80S-IRES complex (Yamamoto et al. 2015; Lancaster et al. 2006) (Supp. Fig. 4h). Most of the 40S-IRES smFRET efficiency distributions were well-fit by single Gaussian functions, irrespective of the IRES labeling position, with a few best fit by a double Gaussian function (Supp. Fig. 4i). However, there were relatively rare transitions from low to high FRET states for single complexes across all conditions (Supp. Fig. 4d-h), some of which notably altered the appearance of the FRET efficiency histogram. As expected, the smFRET efficiency was higher when Cy5-IRES(C44) was the acceptor (∼0.28) in comparison to Cy5-IRES(U56) (∼0.17) (Fig. 3d). In both cases, the mean efficiencies were higher than predicted (Supp. Fig. 4i), likely attributed to the uncertain, dynamic positions of the dyes on the flexible ends of RACK1. Thus, we conclude that RACK1-ybbR is present at its native position on the 40S subunit, and 40S-RACK1-ybbR ribosomes are amenable to single-molecule analysis.

We previously observed conformational dynamics within the 40S-IRES complex at 5 mM Mg^2+^ that were absent at a lower Mg^2+^concentration and in the presence of translation extracts (Fuchs et al. 2015). However, it was unclear whether these measurements were influenced by the bulky tags used for fluorescent labeling: eS25 was C-terminally labeled with a SNAP tag, which is nearly twice its size; and the HCV IRES was labeled at the 5’ end by annealing a 24 nucleotide DNA oligo. We thus completed similar analyses using our improved and much more compact RACK1-IRES smFRET signals for comparison. We observed that the smFRET efficiencies between RACK1 and both IRES constructs were modestly shifted when the Mg^2+^concentration was raised from 2 to 5 mM (Fig. 3d and Supp. Fig. 4). There was a minor population of higher FRET efficiency states (∼0.74) for the Cy5-IRES(C44) acceptor at 5 mM Mg^2+^ that was absent at the lower Mg^2+^concentration. This minor population was also absent in the 80S-IRES complex, which instead had a minor population of even higher FRET (∼1.00). While we currently hesitate to conclude mechanisms from these high FRET states given their rarity and existence at higher-than-physiological Mg^2+^concentrations, the observed changes occur at Mg^2+^ concentrations that are known to promote the transition from the 40S- to 80S-IRES complex (Lancaster et al. 2006), whereby higher Mg^2+^ concentration may be a surrogate for conformational transitions normally induced by protein factors. Collectively, our new labeling strategy revealed similar conformational dynamics as observed previously using bulkier labels, illustrating that the intrinsic flexibility of the 40S-IRES complex is still observed with a compact labeling scheme.

We subsequently probed the association of 40S subunits with the HCV IRES using the RACK1-IRES smFRET signals in real time, and compared the measurements to those made with our prior labeling scheme (Fuchs et al. 2015). We applied a zero-mode waveguide-based (ZMW) system to make real-time smFRET measurements (Chen et al. 2014), and selected the IRES(C44) label since it had higher FRET efficiency with RACK1-ybbR in comparison to IRES(U56). To further improve the signal and ensure an unambiguous binding state, we tested both Cy3 with Cy3.5 as the donor fluorophore. The latter has greater spectral overlap with Cy5 (Buckhout-White et al. 2014) and an increased R_0_, yielding higher FRET intensities at greater distances. We prepared Cy3- and Cy3.5-IRES(C44) RNAs (donors), and 40S-RACK1-ybbR-Cy5 ribosomal subunits (acceptor), and simultaneously immobilized the labeled IRESs within Neutravidin-coated ZMWs on a single chip with 150,000 ZMWs (RS chip) (Fig. 3e). 40S-RACK1-ybbR at 40 nM was added to the surface of the RS chip after the start of data acquisition, and its association with a single immobilized HCV IRES was detected via appearance of smFRET (Fig. 3e and Supp. Fig. 5a). As expected, the Cy3.5-Cy5 pair yielded increased smFRET efficiency (∼0.23) relative to Cy3-Cy5 (∼0.14) (Fig. 4f and Supp. Fig. 5b-d), which was lower than that observed using TIRFM, possibly due to quenching of Cy5 by the aluminum ZMWs (Uemura et al. 2010). 40S association times, defined as the time between the start of the data acquisition and appearance of smFRET, and 40S residence times, defined as the length of the smFRET signal, were nearly identical for both Cy3- and Cy3.5-IRES(C44) molecules (Fig. 4d and Supp. Fig. 5e,f), and similar to our prior measurements of 40S-IRES association kinetics (Fuchs et al. 2015). Thus, by dual-immobilization of Cy3- and Cy3.5-labeled HCV IRESs in ZMWs, we identified an optimal smFRET pair for our single-molecule analyses in ZMWs and validated a strategy that might enable simultaneous tracking of ribosome recruitment to multiple variant RNAs in a single experiment.

**Figure 4.**
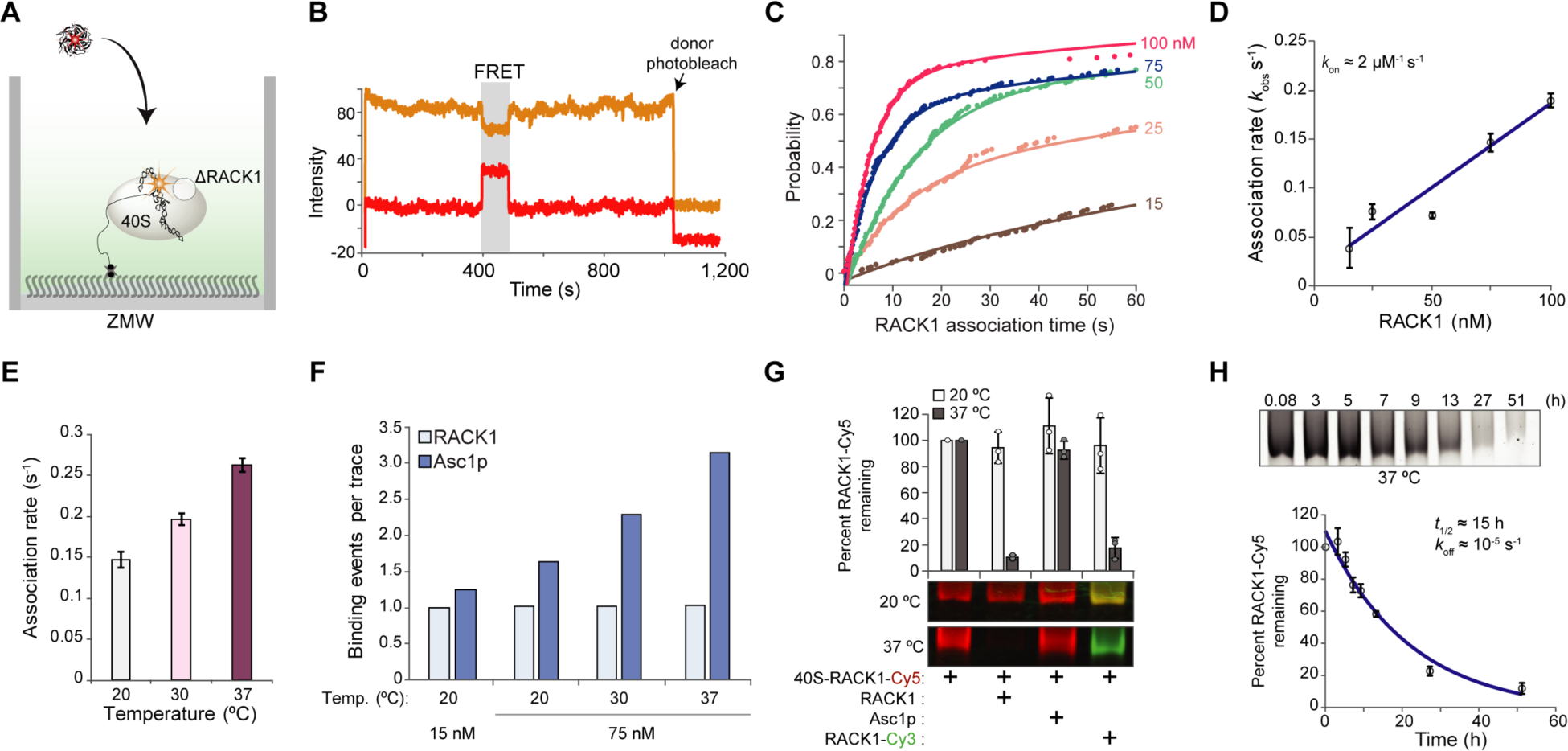
Monitoring RACK1 flux by real-time smFRET in ZMWs. **A.** Schematic of RACK1 delivery to surface-immobilized ΔRACK1 40S in complex with the HCV IRES labeled at C44 with Cy3.5 in ZMWs. **B.** Representative single-molecule fluorescence trace for RACK1-ybbR-Cy5 binding to a single ΔRACK1 40S:Cy3.5-IRES complex indicated by Cy3.5-Cy5 FRET, highlighted in gray. **C.** Cumulative distribution plot of observed RACK1 association times from 0 to 60 seconds. The labels on the right indicate RACK1-ybbR-Cy5 concentration (nM). The lines represent fits to double-exponential functions. **D.** Plot of observed RACK1 association rates (*k_obs_*) of the fast phase at the indicated concentrations. Error bars represent the 95% confidence interval for each value. The line represents a fit to a linear function, with equation y = 0.0017x + 0.0149 and *R^2^* = 0.9225, which yielded the indicated association rate (*k_on_*) for the fast phase. The non-zero intercept may be due to experimental factors such as the presence of unlabeled RACK1-ybbR that competes with the labeled protein (RACK1-ybbR had ∼70% labeling efficiency). **E.** Plot of observed association rates (*k_obs_*) for the fast phase at the indicated temperatures upon delivery of RACK1-ybbR-Cy5 at 75 nM. Error bars represent the 95% confidence interval for each value. **F.** Plot of the observed number of binding events on a single ΔRACK1 40S-IRES complex within a single experiment, indicated by smFRET, for RACK1-ybbR-Cy5 and Asc1p-ybbR-Cy5 delivered at 15 or 75 nM at the indicated temperatures. **G.** Native gel electrophoresis analysis of 40S-RACK1-ybbR-Cy5 at 40 nM following competition for 24 hours with recombinant unlabeled RACK1-ybbR and Asc1p-ybbR at 20-fold excess, and recombinant RACK1-ybbR-Cy3 at 10-fold excess. The gels were analyzed in parallel and are displayed as an overlay of Cy5 (red) and Cy3 (green) fluorescence. The integrated density of each band was quantified using ImageJ, normalized to the respective control sample (lane 1, from left), and the mean (*n* = 3) Cy5 fluorescence was plotted. Error bars represent standard deviation, and circles represent individual data points. **H.** Native gel electrophoresis analysis of 40S-RACK1-ybbR-Cy5 at 40 nM following competition after the indicated times with recombinant unlabeled RACK1-ybbR at 20-fold excess. A representative gel is shown and displays Cy5 fluorescence. Each point on the graph below represents the mean fluorescence intensity remaining relative to the 0.08 h time point, and error bars represent standard deviation. The line represents a fit to an exponential function (*R^2^* = 0.93), which yielded *t_1/2_* = 15 h ± 5 h. The number of traces analyzed (*n*) for panels c, e, and f are listed in Supplementary Figure 9e and 10c.

### RACK1 association with the 40S ribosomal subunit

The above experiments established the functionality of RACK1-ybbR in native 40S ribosomal subunits and an smFRET pair between RACK1 and the HCV IRES suitable for real-time measurements. We next adapted these tools with a reconstituted system to study RACK1 flux on and off the ribosome. After expression in *E. coli*, we purified and fluorescently labeled ybbR-tagged RACK1, as well as mutant versions proposed to have reduced association with the ribosome. These included the D107Y mutation (Kuroha et al. 2010), and a phosphomimetic version (S278E) not expected to interfere with ribosome binding (Jha et al. 2017). Of note, the D107Y mutant was less stable *in vitro* and co-purified with *E. coli* HSP60, suggesting a sub-population may be misfolded or unstable (Supp. Fig. 6). Similarly, a second mutant proposed to disrupt the RACK1-ribosome interaction (R36D/K38E) (Sengupta et al. 2004; Coyle et al. 2009) was also recalcitrant to purification, again likely due to decreased stability (Supp. Fig. 6). These mutants therefore were unsuitable for single-molecule analysis and suggest revisiting interpretations of the many experiments that have used the mutants to deconvolute ribosomal versus extra-ribosomal roles for RACK1, as previously cautioned (Thompson et al. 2016). To circumvent these issues, we hypothesized that the *S. cerevisiae* RACK1 ortholog, Asc1p, would bind the human ribosome with weakened affinity relative to RACK1. Asc1p has multiple substitutions in comparison to RACK1 along the interaction interface with the ribosome (Nielsen et al. 2017) (Supp. Fig. 7), and the human protein associated more weakly than Asc1p to ribosomes in live yeast (Gerbasi et al. 2004). We therefore purified and fluorescently-labeled Asc1p (Supp. Fig. 6), yielding a stable and potentially lower-affinity comparison to wild-type RACK1. Following a similar rationale, we also attempted to purify the RACK1 ortholog from *P. falciparum*, which is absent from structures of its ribosome, but we were unable to purify the protein for similar reasons to the human mutants.

With a small panel of RACK1 proteins predicted to have a range of affinities with the human ribosome in hand, we next tested whether the different RACK1 proteins associated with wild-type and RACK1-deficient 40S subunits *in vitro*. We first purified ΔRACK1 40S ribosomal subunits and determined that they bound fluorescently-labeled HCV IRES with a similar efficiency to wild-type subunits and were competent in Mg^2+^-driven 80S-IRES formation (Supp. Fig 8a,b). These analyses further validated the stability of RACK1-deficient ribosomes and implied that any requirement for RACK1 in HCV IRES activity is independent of mRNA recruitment (Majzoub et al. 2014). Recombinant RACK1 incorporated into ΔRACK1 40S subunits and it did so far more efficiently than to wild-type 40S subunits (Supp. Fig 8c,d). The small amount of RACK1 that incorporated into wild-type ribosomes, which presumably have native RACK1 present, likely reflects a small population that lacked RACK1 or exchange of the native protein with the labeled recombinant version. As hypothesized, the D107Y mutant RACK1 failed to incorporate into WT and RACK1-deficient 40S subunits, while S278E RACK1 incorporated at similar levels as the wild-type protein into both ribosomes (Supp. Fig 8d). While Asc1p incorporated into ΔRACK1 40S subunits, it failed to incorporate into wild-type ribosomes, supporting a lower-affinity than RACK1 for the human ribosome. Furthermore, *in vitro* reconstituted 40S-RACK1-ybbR ribosomal subunits were competent to form 80S complexes on the HCV IRES without notable RACK1 dissociation (Supp. Fig 8e), contrasting a previous structural study of the complex in which ∼50% of particles lacked RACK1 density (Boehringer et al. 2005).

To confirm that recombinant RACK1 incorporates into the native RACK1 binding site, we examined *in vitro* reconstituted 40S ribosomal subunits bound to labeled HCV IRES using single-molecule FRET. In complex with HCV IRES(C44-Cy5), *in vitro* reconstituted 40S-RACK1-ybbR subunits yielded a broad smFRET efficiency distribution with a mean centered at ∼0.31 (Supp. Fig 8f,g). This was remarkably similar to the mean intensity observed with native 40S-RACK1-ybbR subunits (∼0.28) (Supp. Fig. 4i). However, the smFRET efficiency distribution of the reconstituted complex had a broader standard deviation than that of native 40S-RACK1-ybbR (∼0.30 versus ∼0.21), potentially resulting from multiple, heterogeneous FRET states that are difficult to distinguish. These modest differences between native and *in vitro* reconstituted 40S-RACK1-ybbR subunits may represent differences in post-translational modifications of RACK1 (*e.g.*, N-terminal acetylation(Van De Waterbeemd et al. 2018)) or modifications of other ribosomal proteins as discussed above (Supp. Fig. 3d). We also found that human 40S subunits containing Asc1p-ybbR yielded an smFRET distribution similar to both *in vitro* reconstituted and native 40S-RACK1-ybbR subunits, albeit with a small population of high FRET states (mean ∼1.00) (Supp. Fig 8f,g). Of note, following reconstitution, Asc1p had to be included in the TIRFM imaging buffer to facilitate observation of enough smFRET instances for analysis, which was not required for human RACK1 and preliminarily suggested that Asc1p has a shorter residence time on the human 40S subunit. Thus, our findings demonstrate that recombinant RACK1-ybbR incorporated into its native binding site on the ribosome *in vitro* and the reconstituted ribosomes behave similarly to those assembled in cells.

Enabled by our reconstituted system, we next measured the kinetics of RACK1 association to ΔRACK1 ribosomes using real-time smFRET analyses in ZMWs. We immobilized pre-formed complexes of the HCV IRES, labeled at C44 with Cy3.5, and ΔRACK1 40S ribosomal subunits in ZMWs (Fig. 4a). RACK1 association with the 40S-IRES complex was observed via appearance of smFRET (Fig. 4b and Supp. Fig. 9a). As predicted for a single-step bimolecular association, RACK1 displayed concentration-dependent association times (Fig. 4c and Supp. Fig. 9b). After fitting data from the 15, 25, 50, 75, and 100 nM RACK1 deliveries with double-exponential functions, we found that both the rates and amplitudes were concentration dependent (Fig. 4d and Supp. Fig. 9c-e). The fast second-order rate constant was approximately 2 µM^−1^ s^−1^ at 20°C (Fig. 4d), which increased linearly with temperature in a range from 20 to 37°C (see Methods for more details; Fig. 4e and Supp. Fig. 9e-g). The slow second-order rate constant was approximately 20-fold slower (∼0.08 µM^−1^ s^−1^) and plateaued at 30°C (Supp. Fig. 9c-e). The slow phase might represent a sub-population of RACK1-deficient 40S subunits that inhibits RACK1 reincorporation, perhaps due to a post-transcriptional modification on or near the RACK1 binding site, a conformational rearrangement of the 40S-IRES complex, and/or a missing factor that orients the ribosome in a RACK1-accessible conformation. However, the slow phase could instead represent an experimental artifact arising from a source such as non-uniform mixing across the ZMW chip upon delivery of RACK1-ybbR (Chen et al. 2014). Regardless, the fast association rate we observed for RACK1 incorporation is similar to association rates observed for protein-protein (median ≈ 6.6 µM^−1^ s^−1^) and protein-RNA (median ≈ 6.3 µM^−1^ s^−1^) interactions (Schreiber 2002; Gleitsman et al. 2017), and it provides a useful baseline rate for incorporation of a core ribosomal protein into mature human ribosomes.

To test the role of RACK1 sequence on ribosomal association and dissociation kinetics, we probed binding of RACK1 mutants using single-molecule FRET. We first delivered the RACK1 mutant (D107Y) to immobilized 40S-IRES complexes in ZMWs as above. Consistent with a weaker interaction, the D107Y substitution reduced the rate of RACK1 association (Supp. Fig. 9h); however, due to the reduced stability we observed *in vitro*, we suspect that this effect could be due to an overestimation of the folded and active RACK1 concentration. We therefore examined yeast Asc1p association with the human ribosome as a proxy for a stable RACK1 mutant. Upon its delivery to immobilized 40S-IRES complexes, the rate of Asc1p association was dependent on its concentration and temperature (Supp. Fig. 10a-c). The observed fast association rate for Asc1p at 75 nM (∼0.2 s^−1^ at 20°C) was similar to the observed RACK1 association rate (∼0.2 s^−1^) in identical conditions (Supp. Fig. 10b), as estimated via double-exponential functions (Supp. Fig. 9e and 10c). However, in contrast to RACK1, we observed multiple Asc1p binding events separated by as little as 1-3 minutes on a single 40S-IRES complex, which increased with Asc1p concentration and reaction temperature (Fig. 4f and Supp. Fig. 10d-f). Thus, our real-time data suggest that perturbing the RACK1-ribosome interface alters the stability of the interaction.

We next applied single-molecule and bulk approaches to determine the rate of dissociation of wild-type and mutant RACK1 from ribosomes. Wild-type RACK1 and Asc1p residence times on the 40S-IRES complex, defined as the duration of smFRET events, were both concentration-independent (Supp. Fig. 11a-c) and dependent on the power of the excitation laser (Supp. Fig. 11d). Thus, our single-molecule residence time measurements were limited by photobleaching of the dyes, suggesting that they underestimated the lifetime of RACK1-ribosome interactions. To overcome this obstacle, we applied an ensemble approach using native gel electrophoresis. We incubated native 40S-RACK1-ybbR-Cy5 ribosomal subunits with a 20-fold excess of recombinant, unlabeled RACK1-ybbR and quantified the departure of RACK1-ybbR-Cy5 from the ribosome by resolving 40S subunits on a native gel. In this assay, ribosome binding of the unlabeled competitor prevents re-association of the labeled protein upon its dissociation from the ribosome, yielding a time-dependent decrease in fluorescence signal. After 24 hours at 37°C, nearly all of the Cy5 fluorescent signal was lost upon competition with excess RACK1-ybbR (Fig. 4g). Furthermore, there was near-complete exchange of RACK1 upon competition with 10-fold excess RACK1-ybbR-Cy3, indicating that the RACK1 binding site and ribosomal subunits remain intact throughout the experimental window. In contrast, we observed minimal exchange of RACK1 at 20°C or upon competition with 20-fold excess Asc1p at both temperatures. We therefore measured the half-life of RACK1 on the ribosome at 37°C, which we determined to be approximately 15 hours using this bulk exchange assay (Fig. 4h and Supp. Fig. 11e). Together with the observed fast association rate estimated from the 37°C single-molecule experiments, we estimated an equilibrium dissociation constant of ∼10 pM for the RACK1-40S interaction. These results suggest that RACK1 will be stably associated with the ribosome for well over the timeframe of translation in the cell, a process that is typically completed within a few minutes.

## DISCUSSION

Single-molecule fluorescence strategies are powerful approaches to examine the inherent dynamics of protein synthesis. In this work, we establish RACK1 as a new location to label the human ribosome site-specifically with fluorescent dyes via a compact peptide tag and SFP synthase (Yin et al. 2006). By leveraging the proximity of HCV IRES domain II to RACK1 within the 40S-IRES complex, we applied a smFRET signal to examine the kinetics of the RACK1-ribosome interaction. This integration of biochemical, biophysical, and cell-based approaches revealed the remarkable stability of RACK1 on the ribosome. Unlike prokaryotic systems for which ribosome assembly has been reconstituted (Traub and Nomura 1968; Nierhaus and Dohme 1974), such kinetic parameters of RP-ribosome interactions have remained elusive in humans.

Our initial goal was to engineer human ribosomes for improved labeling with fluorescent dyes amenable to single-molecule analysis. RACK1 was an ideal candidate, as it is dispensable for growth in select cell lines, located between the mRNA entry and exit channels, and proximal to binding sites of translation-related factors and viral IRESs. We therefore fused the 11 amino acid ybbR tag onto RACK1, which is among a select group of peptide tags used to label ribosomes in other species (Kaiser et al. 2011; Wang et al. 2012). However, such a short tag has yet to be applied to human ribosomes, and RACK1 is linked to a myriad of translation-related cellular processes. We therefore characterized RACK1-ybbR tagged 40S subunits using orthogonal lines of experiments in cells, and with bulk and single-molecule analyses to ensure their functionality. We found that ribosomes with RACK1-ybbR are functional in cells, biochemically intact, active in simple functional assays, and amenable to single-molecule assays on multiple platforms. In tandem with the segmental-labeling strategy of the IRES, they yielded an unambiguous smFRET signal for the presence of RACK1 within the 40S-IRES complex and enabled us to refine our previous analyses on the dynamics of HCV IRES domain II within the 40S-IRES complex. We also demonstrate their amenability to real-time measurements in ZMWs with complete translation systems. Together, this work highlights the utility of compact labeling schemes for large RNAs and ribonucleoprotein complexes in single-molecule assays. It also illustrates that ybbR and other small peptide tags are prime candidates for tagging human ribosomal proteins at their endogenous genomic loci, expanding the candidates for fluorescent labeling to include essential RPs.

Our *in vitro* experiments collectively demonstrate that RACK1 is a rigid and stable component of the human ribosome. Motivated by the rarity of information the biochemistry of RP-deficient ribosomes and the rates of *in vitro* RP exchange (Kossinova et al. 2008; May et al. 2012), we leveraged the smFRET signal described above with ΔRACK1 40S subunits and RACK1-ybbR to monitor RACK1 binding directly to the ribosome. We showed that RACK1-deficient 40S subunits are functional in simple biochemical assays and nearly identical in composition to wild-type ribosomes. Recombinant RACK1 specifically and unambiguously re-incorporates into human ΔRACK1 ribosomes at the expected location and we observe limited conformational dynamics. The fast second-order rate constant for RACK1 incorporation is approximately 4 µM^−1^ s^−1^ at 37°C, similar to that of Asc1p incorporation (*k_on_* ≈ 2 µM^−1^ s^−1^ at 37°C), which served as our best available proxy for a stable RACK1 mutant. Using a native gel shift assay, we found that RACK1 has a long half-life on the human ribosome *in vitro* (*t_1/2_* ≈ 15 hrs), consistent with the expectation from most structural analyses. Substitutions along the RACK1-ribosome interface, as demonstrated by the use of the distantly-related Asc1p, led to multiple dissociation and re-association events on the 40S subunit over the short time window of our single-molecule experiments. Thus, the high-affinity RACK1-ribosome interaction (*K_D_* ≈ 10 pM) is specified by the slow rate of RACK1 dissociation that is likely mediated by the extensive interface (∼1,800 Å^2^) between RACK1 and the ribosome, which includes contacts with three RPs and 18S rRNA (Nielsen et al. 2017; Sengupta et al. 2004).

Both in our single-molecule assays and in cells, the ribosome is often assumed to have infinite stability over its lifetime. Recent cellular studies have challenged this paradigm and have proposed that variations in the composition of the ribosome can yield specialized functions (Gilbert 2011; Xue and Barna 2012). To overcome the intrinsic stability of the RACK1-ribosome interaction that we observed, however, regulatory control via RACK1 ribosome occupancy would require hours for its stochastic departure from the ribosome or active mechanisms to release RACK1. Indeed, our findings are consistent with the possibility that the ribosome-bound and small pool of free RACK1 can exchange stochastically over the lifetime of a ribosome (*t_1/2_* ≈ 5-7 days) (Dice and Schimke 1972; Nikolov et al. 1987; Defoiche et al. 2009; Mathis et al. 2017). Such exchange has been proposed for other RPs and may take place during viral infections in bacteria (Mizuno et al. 2017) and neuronal functions of mammals (Shigeoka et al. 2018). On the other hand, rapid changes to mature ribosomes could be achieved through the activated release of RACK1, or another RP, perhaps triggered by covalent modifications or other ribosome rearrangements. The ribosomal protein eS10 may be an example of such a phenomenon, as it has been observed at sub-stoichiometric levels in the human ribosome (Shi et al. 2017; Van De Waterbeemd et al. 2018) and is a target for regulatory ubiquitination (Juszkiewicz et al. 2018; Sundaramoorthy et al. 2017). Our *in vitro* findings therefore provide an initial kinetic benchmark for the stability of an integral RP on mature human ribosomes, lending biochemical insight for complementary cellular investigations on the biological consequences and timescales of RP exchange.

The results presented here establish RACK1 as an advantageous location to install a compact fluorescent label on the human ribosome. We demonstrate the utility of this labeling position by defining baseline kinetic parameters for RACK1 association with the ribosome, revealing the remarkable stability of the interaction. We therefore expand the toolkit available for single-molecule studies required to reveal the kinetics that govern human translation and its regulation (Sokabe and Fraser 2018), and provide a biochemical benchmark for the flux of RPs on and off the mature ribosome.

## MATERIALS AND METHODS

### Molecular cloning

To isolate human RACK1 cDNA for cloning, total RNA from HeLa cells was extracted using TRIzol (ThermoFisher, cat.# 15596026)), reverse transcribed and amplified with gene-specific primers, transformed into TOP10 cells, and validated by Sanger sequencing. To make the RACK1-ybbR lentiviral expression construct, RACK1-ybbR was amplified and inserted into pENTR/D-TOPO (ThermoFisher) and then cloned into the lentivirus expression plasmid pLenti CMV Puro DEST (w118-1) (Addgene) by Gateway cloning (ThermoFisher) and validated by sequencing. The open-reading frames (ORFs) for RACK1 proteins were inserted into a pET28c-6xHis-MBP bacterial recombinant expression plasmid backbone downstream of maltose-binding protein (MBP) using a construct derived from a human eIF1A expression construct.(Fraser et al. 2007) The ORF from human RACK1 was subcloned from the lentivirus expression construct, the ORF for *P. falciparum* RACK1 (PfRACK1) was subcloned from *P. falciparum* asexual stage cDNA from the D10 strain (a gift from Michael Boucher, Ellen Yeh lab), and the ORF for *S. cerevisiae* Asc1p was subcloned from a cloned cDNA construct purchased from Genscript (ORF clone OSi04112D). At the 3’ end of each construct was sequence encoding for the ybbR peptide tag and a stop codon (GATTCTCTTGAATTTATTGCTAGTAAGCTTGCGTAG). We refer to the proteins encoded by these plasmids as 6xHis-MBP-RACK1/Asc1p-ybbR fusion proteins. The D107Y, R38D/K40E, and S278E amino acid substitutions were introduced into human RACK1 using the QuickChange II XL site-directed mutagenesis kit (Roche). All plasmids are available upon request.

### Cell growth

HAP1 cells were grown at 37°C with 5% CO_2_ in Iscove’s Modified Dulbecco’s medium (IMDM) (Gibco, cat.# sh30228) supplemented with 10% (v/v) fetal bovine serum, 2 mM L-glutamine (Gibco), and 1X penicillin/streptomycin (Gibco). As indicated, wild-type and ΔRACK1 (clone E3-A5) HAP1 cells were used in our studies (Carette et al. 2011; Jha et al. 2017).

### Lentiviral transductions

Lentivirus packaging was conducted either at the Neuroscience Gene Vector and Virus Core of Stanford University or performed in-house using pLenti constructs by co-transfection with ΔVPR, VSV-G, and pAdVAntage packaging plasmids into HEK293FT cells using FuGENE HD (Promega) (Campeau et al. 2009). Cells were transduced with lentivirus-containing media 24 hours after seeding in 6-well plates at 250,000 cells per well. Lentivirus-containing media was replaced with fresh media after 24 hours, and after an additional 24 hours, transduced cells were selected by the addition of 1 ug/mL puromycin. Following five days under puromycin selection, surviving cells were expanded in culture for characterization and ribosome purifications.

### Polysome profiling

HAP1 cells were grown by routine passaging and cells were brought to ∼80% confluence in 150-mm culture dishes, receiving fresh IMDM media 4-6 hours prior to harvesting. Immediately before harvest, 100 μg/ml cycloheximide (CHX) was added to media, and cells were incubated for 3 minutes at 37°C. Media was removed by aspiration, adherent cells were washed twice with ice-cold PBS (containing 100 μg/ml CHX). PBS was aspirated from cells, which were subsequently dissociated from plates by scraping and transferred to Eppendorf tubes in residual PBS. An equal volume of 2X cell lysis buffer (30 mM Tris-HCl pH 7.5, 150 mM NaCl, 2% Triton X-100, 10 mM MgCl_2_, 2 mg/mL heparin, 0.2 mg/mL CHX, and 2 mM DTT) was mixed with cell suspensions, and lysis proceeded for 10 minutes on ice. Lysate was clarified to remove nuclei at 8,400 x *g* for 5 min at 4°C with a microcentrifuge, and lysates were measured with a nanodrop to normalize gradient inputs by absorbance at 260 nm. Using 14×89 mm thin-wall polypropylene tubes (Beckman Coulter ref. 331372), 10-60% sucrose gradients were prepared in 20 mM Tris-HCl pH 7.5, 150 mM NaCl, 5 mM MgCl2, 0.1 mg/mL CHX, and 1 mM DTT, and linearized using a BioComp 107 Gradient Master. Post-nuclear lysate samples from a single 150-mm dish (500 uL of ∼1000 ng/uL RNA) was loaded onto each pre-chilled sucrose gradient and centrifuged at 150,000 x *g* (35,000 rpm) using a SW41 Ti rotor for 2 hours 45 min at 4°C. Gradients were fractionated with a Brandel gradient fractionation system using a 5 mm UV cell detector, pumping at 0.75 mL per minute, and collecting fractions across the length of the gradient. Protein was extracted from each 0.75 mL fractions by methanol extraction. Briefly, 0.1 mL of each fraction was mixed with 0.4 mL methanol, vortexed, and spun for 10 sec at 9,000 x g in a microcentrifuge. To each sample, 0.1 mL chloroform was added, followed by additional vortexing. Next, 0.3 mL nanopore H_2_0 was added to each fraction with vortexing for phase separation. The upper phase was discarded, and 0.3 mL was added to the lower phase with more vortexing. The protein was then pelleted by centrifuging at max speed in a microcentrifuge for 2 minutes, from which the supernatant was removed, and the pellet was further dried using a SpeedVac. Dried protein pellets were resuspended in 0.01 mL 8 M Urea/ 100 mM Tris-HCl pH 8.0. Solubilized proteins were then used for SDS-PAGE and western blotting.

### Proliferation assay

Proliferation assays were performed as described (Johnson et al. 2018). Briefly, wild-type, ΔRACK1, and ΔRACK1+RACK1-ybbR HAP1 cells were dissociated from culture plates with trypsin and counted for live cells by trypan blue. Cells were then seeded into 96-well plates at 10,000 live cells/well with 10 replicates per cell line, assaying over 4 days of growth using the Vybrant MTT Cell Proliferation Assay Kit (ThermoFisher). Absorbance values at 570 nm were determined using a Synergy Neo2 instrument (BioTek).

### Immunoblotting

Western blotting was performed as described (Johnson et al. 2018). When re-probing blots, HRP-conjugated secondary antibodies were either inactivated by incubation with sodium azide in 5% skim milk/TBST or stripped with Restore Stripping Buffer (ThermoFisher). The following antibodies were used in this study: anti-RACK1 (Cell Signaling, ref. 4716S, 1:1000), anti-RACK1 (Santa Cruz, ref. 17754, 1:500), RPL5 (Genetex, ref. 101821, 1:1000), uS10/RPS20 (abcam, ref. 133776, 1:1000), eS10/RPS10 (Genetex, ref. 101836, 1:1000), uS5/RPS2 (Santa Cruz, ref. 130399, 1:500), and uS3/RPS3 (Bethyl, ref. 303-840A, 1:1000).

### Ribosome purifications

40S and 60S ribosomal subunits from wild-type, ΔRACK1, and ΔRACK1+RACK1-ybbR HAP1 cells were purified as described with a few optimizations (Fuchs et al. 2015). 20-40 x 150-mm culture dishes were grown to ∼80% confluency. Media was removed, cells were washed with ice-cold PBS, and PBS was removed via aspiration. In the residual PBS, cells were scraped to release them from the dish, combined into a 15 mL culture tube, and pelleted by centrifugation at 1,000 x *g* for 10 min in an Eppendorf 5810 R centrifuge equipped with a swinging bucket rotor. Cell pellets were resuspended in ice-cold PBS and re-pelleted via centrifugation to wash. PBS was removed via aspiration, and cell pellets were resuspended in an equal volume of fresh, ice-cold PBS (typically 1-2 mL). An equal volume of 2X lysis buffer (30 mM Tris-HCl pH 7.5, 300 mM NaCl, 20 mM MgCl_2_, 2% (v/v) Triton-X 100, 4 mM DTT, 2 mg/mL heparin, and 1X CoMplete Mini protease inhibitor cocktail (Roche)) was added, cells were mixed gently by multiple inversions of the culture tube, and cells were incubated on ice for 10 min to complete lysis. Cell lysates were cleared of debris and nuclei via centrifugation at 42,000 x *g* (10,000 rpm) at 4°C for 10 min in a Fiberlite F21 rotor (ThermoFisher, cat.# 46923). The supernatant was layered directly onto a high-salt sucrose cushion (20 mM Tris-HCl pH 7.5, 500 mM KCl, 30% (v/v) sucrose, 10 mM MgCl_2_. 0.1 mM EDTA pH 8.0, and 2 mM DTT). Ribosomes were pelleted via centrifugation at 63,000 x *g* for 16-18 hours at 4°C using a Type 80 Ti rotor (Beckman Coulter). Ribosome pellets were dissolved in resuspension buffer (20 mM Tris-HCl pH 7.5, 500 mM KCl, 7.5% (v/v) sucrose, 2 mM MgCl_2_, 75 mM NH_4_Cl, 2 mM puromycin, and 2 mM DTT) by gentle pipetting. To complete resolubilization and splitting of the ribosomal subunits, ribosomes were incubated at 4°C for 1 hour followed by an incubation at 37°C for 1.5 hours. To isolate 40S and 60S ribosomal subunits, the solution was layered directly onto a pre-chilled linear 10-30% sucrose gradient (with 20 mM Tris-HCl pH 7.5, 500 mM KCl, 6 mM MgCl_2_) in 25×89 mm polypropylene centrifuge tubes (Beckman Coulter, ref. 326823) prepared using the BioComp 107 Gradient Master. Gradients were centrifuged at 49,123 x g for 16 hours at 4°C using an SW32Ti rotor (Beckman Coulter, ref. 369694). Gradients were fractionated with a Brandel gradient fractionation system using a 1 mm UV cell detector, pumping at 1.5 mL per minute, and collecting 0.75 mL fractions across the length of the gradient. Fractions containing 40S and 60S ribosomal subunits as identified via absorption at 254 nM were pelleted via centrifugation at 63,000 x *g* for 20 hours at 4°C using a Type 80 Ti rotor (Beckman Coulter). Ribosomal subunits were resuspended in storage buffer (30 mM HEPES-KOH pH 7.4, 100 mM KOAc, 5 mM Mg(OAc)_2_, 6% (v/v) sucrose, and 2 mM DTT), aliquoted, flash frozen in liquid N_2_, and stored at −80°C. Ribosomal subunits were characterized by SDS-PAGE (BioRad Any kD precast gels), the Pierce Silver Stain Kit (ThermoFisher), western blotting, and mass-mapping with the Stanford PAN facility.

### SFP synthase labeling of ybbR peptides

SFP synthase was expressed recombinantly in bacteria using the pET29-Sfp construct (a gift from Jun Yin at Georgia State University) (Yin et al. 2006). Protein expression was performed as described, and protein was subsequently used to catalyze ybbR labeling reactions. Sulfo-Cyanine3 (cat. # 11380) and Sulfo-Cyanine5-maleimide (cat. # 13380) was purchased from Lumiprobe. Coenzyme A trilithium salt (CoA) was purchased from Sigma-Aldrich (cat.# C3019). CoA was reacted with maleimide-dyes as described and CoA-dyes were purified using a 5-50% gradient of acetonitrile in water with 0.1% TFA using an HPLC equipped with a C18 column.(Yin et al. 2006) Successful synthesis and purification of CoA-dye products was verified by LC-MS analysis. After confirming the published synthesis and purification scheme, we instead used the crude CoA-dye reactions without HPLC purification, by quenching excess maleimide with DTT and directly labeling ybbR-tagged proteins with SFP synthase.

40S ribosomal subunits containing RACK1-ybbR were fluorescently labeled with Cy3 or Cy5 by incubating 40S subunits with 2-4 molar excess SFP synthase and CoA-Cyanine dyes at 37°C for 2 hours in ybbR-labeling buffer (50 mM HEPES-KOH pH 7.5, 100 mM NaCl, 10 mM MgCl_2_, 10% (v/v) glycerol, and 1 mM DTT). To remove free dye and SFP synthase, labeled 40S subunits were layered onto a low-salt sucrose cushion (30 mM HEPES-KOH pH 7.5, 100 mM KOAc, 0.5 M sucrose, 5 mM Mg(OAc)_2_, and 2 mM DTT), and centrifuged at 287,582 x *g* (90,000 rpm) for 1 hr at 4°C in a TLA100.2 rotor (Beckman, ref. 362046) in 11×34 mm thick-wall polycarbonate ultracentrifuge tubes (Beckman, ref. 343788). Ribosome pellets were washed once and subsequently resuspended with ribosome storage buffer, aliquoted, flash frozen in liquid N_2_, and stored at −80°C. Recombinant proteins were labeled with cyanine dyes by incubating 10 µM 6xHis-MBP-RACK1-ybbR proteins with 2 µM SFP synthase and 15 µM CoA-Cy3/5 dyes at 37°C for 2 hours. Free dye was removed via purification over 10DG-desalting columns (Bio-Rad, cat.# 7322010) equilibrated in TEV buffer (20 mM Tris-HCl pH 8.0, 150 mM NaCl, 10% (v/v) glycerol, 10 mM imidazole, and 5 mM β-mercaptoethanol). Dye-labeled proteins and ribosomes were characterized as indicated by SDS-PAGE (BioRad Any kD precast gels), fluorescent scanning using a Typhoon instrument, the Pierce Silver Stain Kit (ThermoFisher), western blotting, and mass-mapping with the Stanford PAN facility.

### RACK1 recombinant protein expression and purification

Recombinant 6xHis-MBP-RACK1-ybbR fusion proteins were expressed and purified from *E. coli* BL21(DE3) cells. Two to four liters of cells were grown at 37°C in LB supplemented with 50 µg/mL kanamycin to OD_600_ of 0.5, and cells were rapidly cooled on ice. Expression of the fusion proteins was induced by addition of 0.5 mM IPTG, and cells were maintained in a shaking incubator for 16 hours at 17°C. Cells were harvested by centrifugation at 5,000 x *g* for 10 min at 4°C in a Fiberlite F9 rotor (ThermoFisher, cat.# 13456093).

Cells were lysed by sonication in lysis buffer (20 mM Tris-HCl pH 8.0, 300 mM NaCl, 10% (v/v) glycerol, 10 mM imidazole, and 5 mM β-mercaptoethanol), and lysates were cleared by centrifugation at 38,000 x *g* for 10 min at 4°C in a Fiberlite F21 rotor followed by filtration through a 0.22 µm syringe filter. Clarified lysate was loaded onto a Ni-NTA gravity flow column equilibrated in lysis buffer, washed with 20 column volumes (CV) of lysis buffer, 20 CV of wash buffer (20 mM Tris-HCl pH 8.0, 1 M NaCl, 10% (v/v) glycerol, 25 mM imidazole, and 5 mM β-mercaptoethanol), and 10 CV of lysis buffer. Recombinant proteins were eluted with six sequential CV of elution buffer (20 mM Tris-HCl pH 8.0, 300 mM NaCl, 10% (v/v) glycerol, 250 mM imidazole, and 5 mM β-mercaptoethanol), and fractions with recombinant protein as identified by SDS-PAGE analyses were dialyzed overnight at 4°C into ybbR-labeling buffer (50 mM HEPES-KOH pH 7.5, 100 mM NaCl, 10 mM MgCl_2_, 10% (v/v) glycerol, and 1 mM DTT) or TEV Buffer (20 mM Tris-HCl pH 8.0, 150 mM NaCl, 10% (v/v) glycerol, 10 mM imidazole, and 5 mM β-mercaptoethanol).

Fluorescently-labeled and unlabeled proteins were dialyzed overnight in the dark at 4°C into TEV buffer in the presence of 1-1.5 mg of TEV protease. To separate 6xHis-tagged proteins from RACK1-ybbR(Cy3/5), the proteins were passed through a second Ni-NTA gravity column equilibrated in TEV buffer to capture 6xHis-MBP, TEV protease, and SFP synthase. Flow-through containing RACK1-ybbR(Cy3/5) was further purified using size exclusion chromatography using a Superdex 75 column (23 mL) equilibrated in SEC buffer (20 mM HEPES-KOH pH 7.5, 150 mM KOAc, 10% (v/v) glycerol, and 1 mM DTT). Fractions containing RACK1-ybbR(Cy3/5) were concentrated using a 10 kD MWCO Amicon Ultra centrifugal filter, aliquoted, flash frozen on liquid N_2_, and stored at −80°C. The concentration was determined via absorption at 280 nm using a nanodrop for total protein, and at 548 nM or 646 nM for Cy3 or Cy5 labeled proteins, respectively. A concentration range of indicated recombinant proteins was analyzed by thermal melt assays in SEC buffer using SYPRO Orange (Molecular Probes, cat.# S6651) using a Biorad CFX96 Touch Real-Time PCR C1000 System (ramping from 4 to 100°C at 1°C per minute).

### RNA transcription and fluorescent labelling

For single-molecule experiments, HCV IRES RNA was produced and labeled with cyanine dyes as described (Johnson et al. 2018). Briefly, an acceptor RNA was transcribed *in vitro* by T7 with GMP-priming for segmental labeling. Synthetic RNA donor oligos with cyanine dyes installed terminally at C44 or internally at U56 (TriLink) were ligated to the acceptor RNA using T4 RNA ligase 1 (NEB) (Fig. 3 and Supp. Figure 4). For TIRFM experiments, RNAs were labeled at C44 or U56 with Cyanine5, and for ZMW experiments, RNAs were labeled at C44 with Cyanine3 or Cyanine3.5.

For gel shift experiments, HCV IRES RNA that lacked the 3’ extension used in single-molecule experiments was transcribed *in vitro* using T7 polymerase, the RNA was extracted using phenol:chloroform:isoamyl alcohol, ethanol precipitated, and purified via size-exclusion chromatography on a Superdex 200 column (23 mL) equilibrated with 10 mM Bis-Tris pH 7 and 100 mM NaCl. RNA was concentrated using a 10 kD MWCO Amicon Ultra centrifugal filter, aliquoted, flash frozen on liquid N_2_, and stored at −80°C. The concentration was determined via absorption at 260 nm using a nanodrop.

To end label RNA with Cy5 dye, purified RNA was incubated in 100 mM NaOAc pH 5.5 and 5 mM KIO_4_ for 30 minutes on ice in the dark. The reaction was quenched by addition of 10 mM ethylene glycol and incubation for 5 min on ice in the dark. Oxidized RNA was ethanol precipitated, resuspended in 100 mM NaOAc, and incubated with 5-10 fold molar excess Cy5-hydrazide (Lumiprobe, cat. # 13070) for 6 hours at room temperature in the dark. Free dye was removed via purification over 10DG-desalting columns (Bio-Rad, cat.# 7322010). Full-length, end-labeled RNA was purified by extraction from acrylamide gels following electrophoresis, ethanol precipitated, aliquoted, flash frozen, and stored at −80°C. The dye concentration was determined via absorption at 650 nm using a nanodrop spectrophotometer.

### Refolding of HCV IRES

HCV IRES was diluted to 0.5 to 1 µM in refolding buffer (20 mM cacodylate-NaOH pH 7.0, 100 mM KCl, and 1 mM EDTA pH 8), heated to 95°C for 2 minutes, and slowly cooled to room temperature (over ∼ 45 min). For single-molecule experiments, a biotinylated DNA oligo that anneals to the 3’ end of the RNA on an artificial extension downstream of the IRES was included at 2-fold molar excess. Once cooled, 4 mM MgCl_2_ was added to quench the EDTA.

### Native gel electrophoresis

Ribosomes, and complexes of ribosomes and HCV IRES were analyzed using acrylamide/agarose composite gels as described (Johnson et al. 2018). Briefly, 2.75% acrylamide (37.5:1), 0.5% Nusieve GTG agarose composite gels were prepared in the following buffer: 25 mM Tris-OAc pH 7.5, 4 mM KOAc, 2 mM Mg(OAc)_2_, 0.5 mM DTT, 2.5% glycerol, 0.1% (v/v) TEMED, and 0.1% (v/v) fresh ammonium persulfate. The gels were cooled at 4°C for 20 minutes and allowed to further polymerize at room temperature for 2 hours. Gels were pre-run in ice-cold running buffer (25 mM Tris-OAc pH 7.5, 4 mM KOAc, and 2 mM Mg(OAC)_2_) for 30-60 min at 4°C. All native gel electrophoresis assays were repeated at least three times, and respective ribosome, protein, and RNA samples were derived from a single purification. For gel shift assays with HCV IRES, the indicated amounts of purified ribosomal subunits were incubated for 10 min at 37°C with RNA in ribosome assay buffer (30 nM Cy5-labeled IRES in 30 mM HEPES-KOH pH 7.4, 100 mM KOAc, and 2 mM Mg(OAc)_2_). For analysis of recombinant protein incorporation into purified ribosomal subunits, 40 nM of purified recombinant proteins labeled with Cy5 were incubated for 10 min at 37°C with the indicated amount (titration experiment) or 75 nM (single concentration experiment) of 40S ribosomal subunits in ribosome assay buffer. For analysis of 80S ribosome complex formation, 60 nM of the indicated 40S and 60S ribosomal subunits, and 30 nM of Cy5-labeled IRES were incubated for 10 min at 37°C in ribosome assay buffer with the indicated amount of MgCl_2_.

### Single-molecule spectroscopy

For equilibrium experiments, 100 nM refolded HCV IRES annealed to a biotinylated DNA oligo was incubated with 200 nM of 40S(RACK1-ybbR-Cy3) ribosomal subunits for 10 min at 37°C in ribosome assay buffer. Conditions were identical for experiments with 80S ribosomes except that 400 nM 60S ribosomal subunit was included and ribosome assay buffer was supplemented with 3 mM Mg(OAc)_2_ (5 mM total). For experiments with *in vitro* reconstituted ribosomes, 1 µM of ΔRACK1 40S ribosomal subunits were first incubated with 2 µM of Cy3-labeled recombinant protein for 10 min at 37°C in ribosome assay buffer, and subsequently, 200 nM of in vitro reconstituted 40S ribosomal subunits were incubated with refolded HCV IRES for 10 min at 37°C in ribosome assay buffer. Ribosome-IRES complexes were immobilized on a channel of neutravidin-coated quartz slides by addition of 40 µL 400 pM complex (based on the IRES concentration) and incubating for 5 min. Unbound complexes were washed from the channel using ribosome assay buffer supplemented with 2 mM Trolox (TSY), 2 mM protocatechuic acid (PCA), and 0.06 U/µL protocatechuate-3,4-dioxygenase (PCD). Complexes were imaged on a prism-based total internal reflection fluorescence (TIRF) microscope as described(Aitken et al. 2008), using 532 and 647 nm excitation lasers with movies collected at 5 frames per second and EM gain set to 650. Short movies (50 frames) were collected using either (single) or both (dual) excitation lasers, while long movies (1,500 frames) were collected using excitation with the 532 nm laser. Colocalized molecules were identified and analyzed using custom MATLAB (R2017a, MathWorks) scripts (available upon request).

For real-time experiments in zero-mode waveguides (ZMWs), all imaging was conducted using a modified Pacific Biosystems RS II DNA sequencer as described (Chen et al. 2014; Johnson et al. 2018). ZMW chips purchased from Pacific Biosciences were initially washed with TP50 buffer (50 mM Tris-OAc pH 7.5, 100 mM KCl), coated with neutravidin by incubating 20 µL of 75 nM neutravidin in TP50 supplemented with blocking oligo for 5 minutes, and washed 3-6X with ribosome assay buffer. 40S-IRES complexes were formed by incubating 1 µM ΔRACK1 40S ribosomal subunits with 200 nM refolded HCV IRES labeled at C44 with Cy3.5 annealed to a biotinylated DNA oligo. Complexes were immobilized on the neutravidin-coated ZMW surface by incubating 40 µL of 1 nM of complex (via the IRES) for 5 minutes. Unbound complex was removed via 3-5 washes with ribosome assay buffer. Immediately prior to imaging, buffer was removed from the surface of the ZMW chip and replaced with 20 µL ribosome assay buffer supplemented with 2 mM TSY, 2 mM PCA, and 0.06 U/µL PCD. Upon start of image acquisition, 20 µL of Cy5-labeled recombinant protein in the same buffer was delivered to the chip surface. For experiments with delivery of 40 nM 40S-RACK1-ybbR-Cy5, all conditions were the same except 40 µL of 0.5 nM HCV IRES labeled with either Cy3 or Cy3.5 at C44 was immobilized on the ZMW surface. All movies were collected at 5 frames per second for 20 minutes to maximize the capture of single-molecule photobleaching events of the Cy3.5 dye. Unless noted, the 532 nm laser was used at 0.8 µW/ µm^2^ illumination and the temperature of the imaging chamber was 20°C. The “low laser power” experiment was conducted using the 532 nm laser at 0.32 µW/ µm^2^ illumination at 20°C. All movies were analyzed using custom MATLAB scripts (available upon request) to extract fluorescence intensity and kinetic parameters. The respective ribosome, protein, and RNA samples were derived from a single purification for all equilibrium and real-time single-molecule analyses.

For the single-molecule analyses, *n* represents the number of molecules that were analyzed, unless otherwise noted. For robust kinetic fits, we typically analyzed >175 molecules per condition, and only wells with a single IRES molecule, identified by a single-step photobleaching event of the donor fluorophore. To overcome non-uniform mixing on the surface of the ZMW chip, which occurs within 1 s at the center and is complete at ∼7 s, we concentrated our analyses on molecules present in the center of the chip (Chen et al. 2014). At the lowest concentration (5 nM), the slow rate was dominant, and we were unable to confidently fit the association times with single or double exponential functions. For all analyses that compare smFRET efficiencies, smFRET events were assigned to a single state and single molecules were identified by a single-step photobleaching event of the donor fluorophore, based on relative fluorescence intensities. We then corrected the background fluorescence present in each optical channel and calculated distributions of observed smFRET efficiencies for the respective dye pairs.

The 30°C and 37°C real-time association experiments were conducted with the PacBio RS II imaging chamber set to 28°C and 35°C, respectively, to account for the increased temperature due to laser excitation. We delivered RACK1 at 75 nM for several reasons. First, the fast association rate was dominant, which enabled robust fitting of the data. Second, evaporation of the sample is a concern at the highest temperature in our microscope. We thus focused our measurements on the impact of temperature on fast binding events, which predominately occurred within 30 seconds. Third, this concentration was within a window that would allow observation of faster or slower binding events.

## Data and code availability

The single-molecule datasets generated during and/or analysed during the current study are available from the corresponding author upon request.

All custom MatLab analysis scripts are available upon request or are available on GitHub (https://github.com/corsepius/FRETAutomation).

## ACKNOWLEDGEMENTS

We are grateful to Alexey Petrov, Karim Majzoub, and Jan Carette’s lab for helpful discussions. We thank Peter Sarnow and his lab for sharing cell culture equipment, and Miguel Mata for experimental advice on polysome profiling. We thank Oli Duss for helpful discussions and critical reading of the manuscript. Josh Yim and Matt Bogyo’s lab generously shared equipment and knowledge for use of their HPLC for CoA-dye purification. Ethan LaFontaine helped with cloning of RACK1-ybbR into a lentiviral plasmid. The Stanford PAN facility provided support through mass mapping analysis, and thermal melt experiments were conducted at the ChEM-H MSKC. A.G.J. was supported by a National Science Foundation Graduate Research Fellowship (DGE-114747); C.P.L. is a Damon Runyon Fellow supported by the Damon Runyon Cancer Research Foundation (DRG-#2321-18); J.W. is supported by a postdoctoral scholarship from the Knut and Alice Wallenberg Foundation; N.C.C. is supported by a R35 grant from the NIH in Michael Levitt’s lab (GM122543); J.C. was supported by a Stanford Bio-X fellowship; and G.F. is supported by the University at Albany Faculty Research Awards Program (FRAP). Research on eukaryotic translation in the laboratory of J.D.P. is funded by the National Institutes of Health (AI047365, AI099506 and GM113078).

---

**Supplementary Figure 1.**
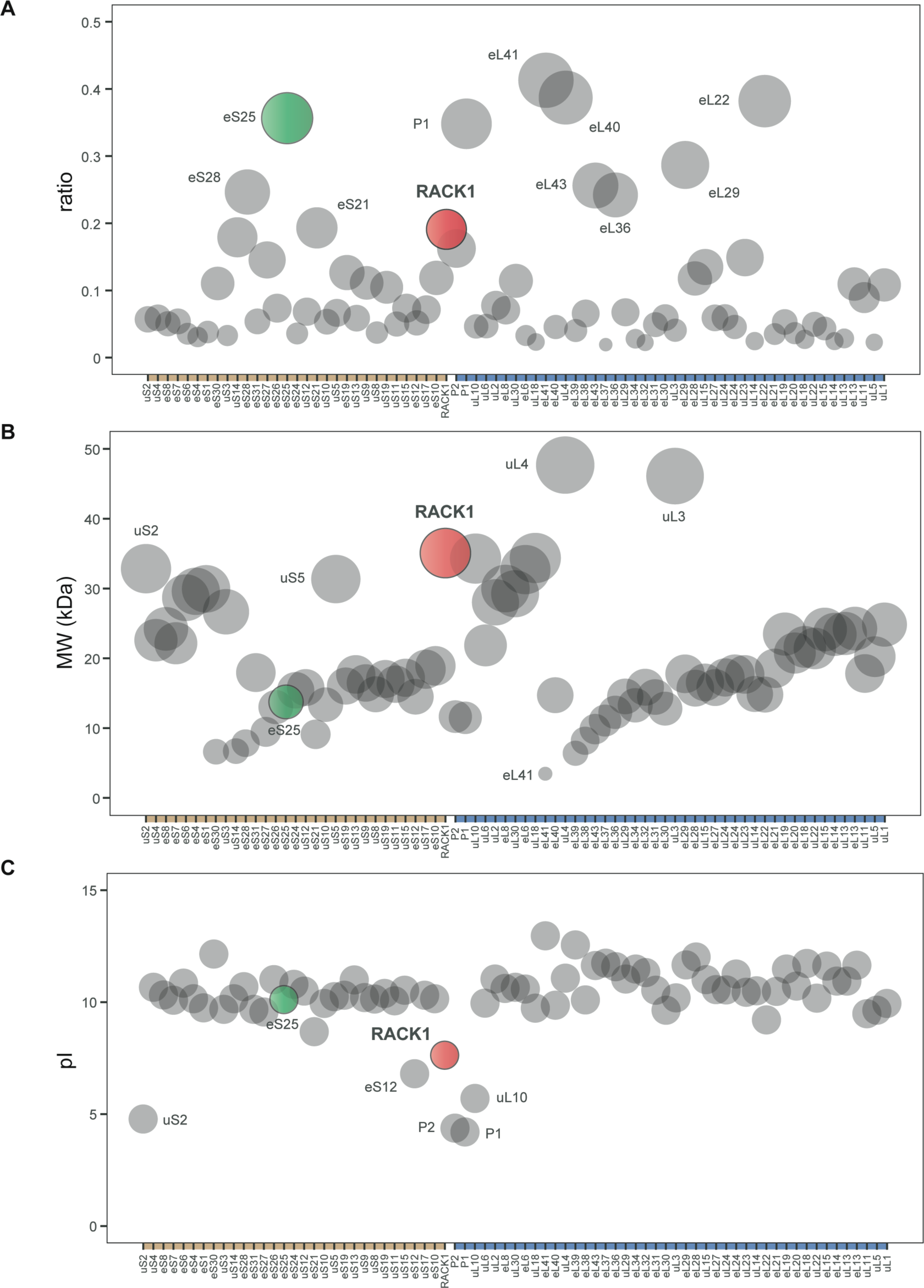
Properties of human ribosomal proteins. For all plots, the small subunit ribosomal proteins are plotted on the left, while those of the large subunit are on the right (tan and blue regions of x-axis, respectively). Proteins are annotated with the new nomenclature for ribosomal proteins, and ribosomal protein paralogs were omitted in this analysis. RACK1 is indicated in red and eS25 is indicated in green. For a and b, the size of each plotted circle reflects the magnitude of the y-axis value. **A.** Ratio of sense to antisense gene-trap insertions from essentiality screen in HAP1 cells (Blomen et al. 2015). While the absolute number of insertions can limit confidence in assigning essential genes, this metric allows an approximation for RP genes that may be deleted without substantial fitness costs (higher ratio ∼ less essential). This is supported by our ability to generate isogenic knockouts of RPS25 and RACK1, but the essentiality of several of the other high ratio genes have yet to be rigorously analyzed in human cells. **B.** Size distribution of human ribosomal proteins based on their predicted molecular weight (MW) in Daltons (Da). **C.** Isoelectric point (pI) distribution of human ribosomal proteins. For b and c, protein statistics were retrieved from Uniprot.

**Supplementary Figure 2 (related to Figure 1).**
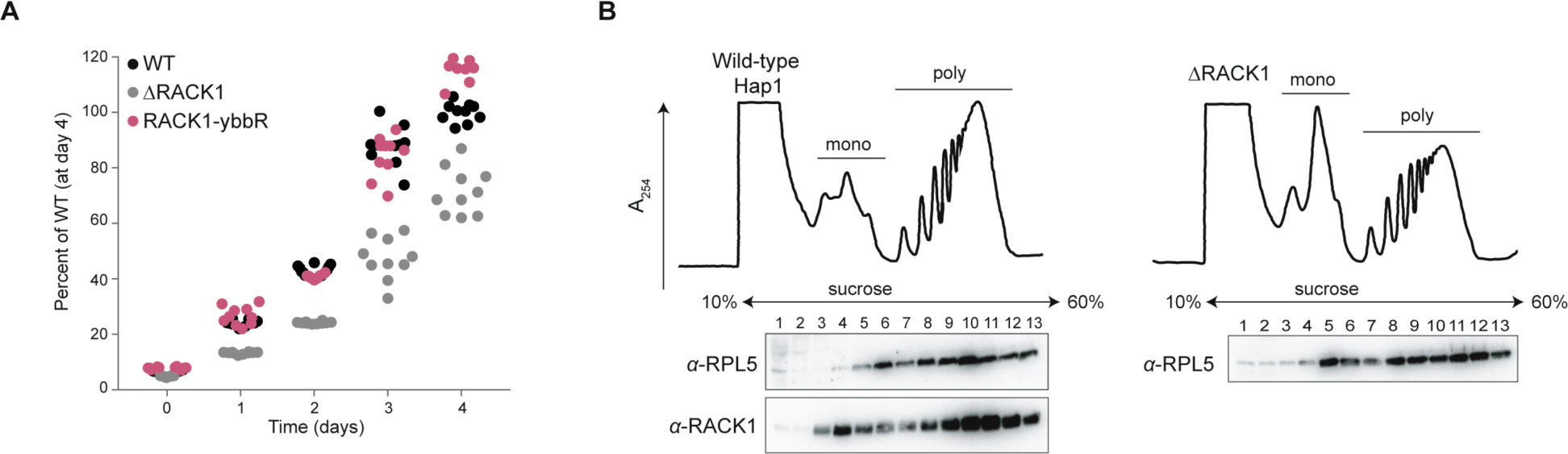
RACK1-ybbR incorporates into translating ribosomes in cells. **A.** Plot of the proliferation of the indicated cell lines as measured by MTT assay relative to the wild-type cells at day 4. Each circle represents cells from a distinct well in a 96-well plate, and these data were used to generate the line graph in Figure 1b. **B.** UV absorbance traces (at 254 nm) of polysome profiling and associated western blot analyses. “Mono” and “poly” refer to peaks that correspond to monosomes and polysomes, respectively.

**Supplementary Figure 3 (related to Figure 2).**
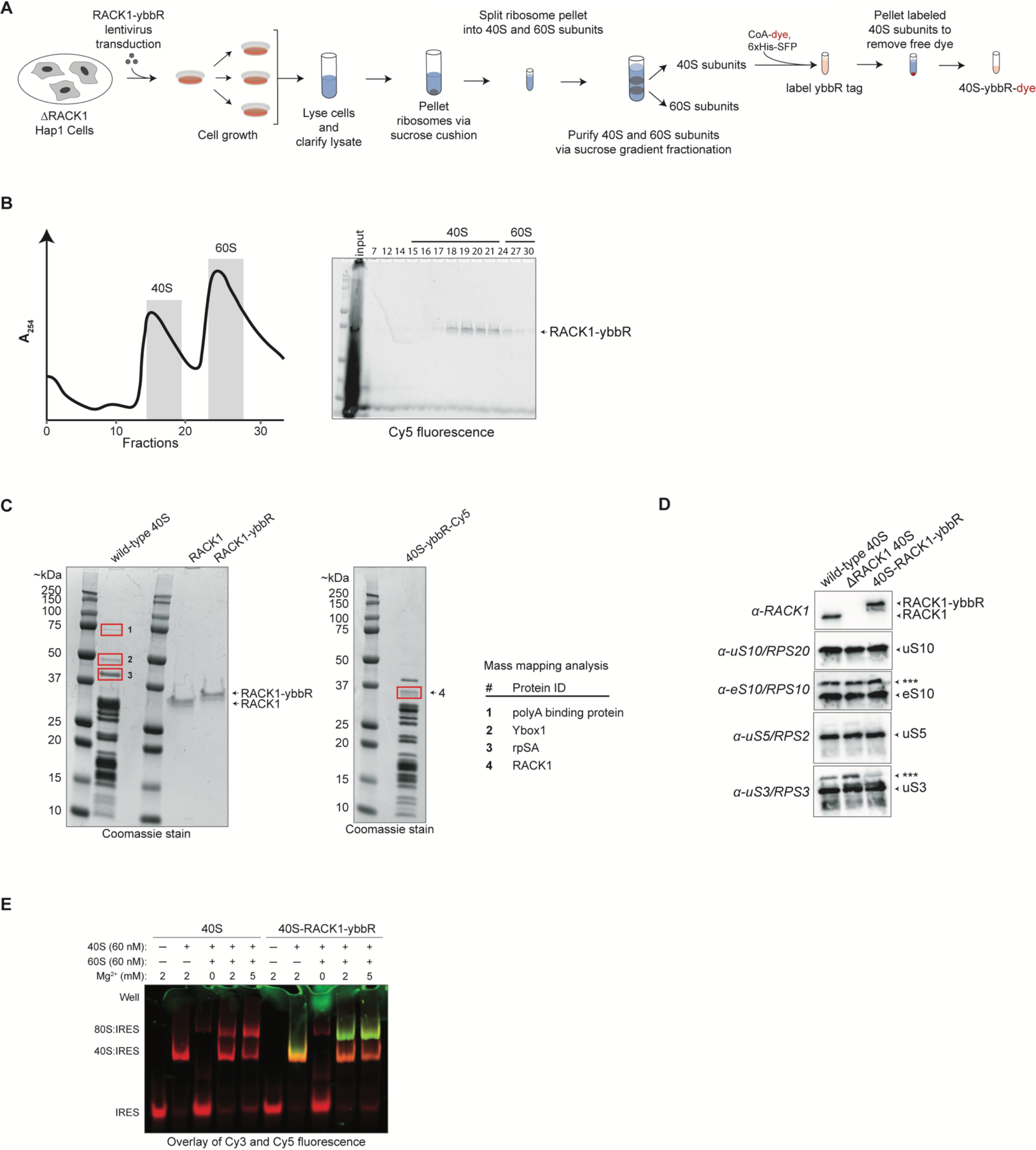
Purification and biochemical characterization of ribosomes labeled with RACK1-ybbR. **A.** Strategy to purify and fluorescently label ribosomal subunits from the human HAP1 cell line. **B.** A representative UV absorbance trace (at 254 nm) (left) and an image of a gel scanned for Cy5 fluorescence (right) following SDS-PAGE analysis of the sucrose gradient fractionation step from our initial attempt to fluorescently-label 40S-RACK1-ybbR subunits. Crude 80S ribosomes (after the first sucrose cushion) from RACK1-ybbR expressing cells were resuspended in a standard high-salt and puromycin-containing subunit splitting buffer supplemented with 10 mM Mg^2+^. After incubation with SFP synthase and CoA-Cy5, we quenched the excess magnesium, split the ribosomes into individual subunits, and purified them using a 10-30% sucrose gradient followed by fractionation, with each fraction examined for Cy5 fluorescence using SDS-PAGE. The single band of Cy5 fluorescence matched the molecular weight of RACK1-ybbR and was primarily localized to the 40S ribosomal subunit peak. Despite its initial promise, however, these ribosomes were poorly labeled, which we attributed to the high concentration of monovalent salt and other contaminants present in the crude 80S ribosomes during labeling. Nevertheless, the trace represents the typical separation of 40S and 60S ribosomal subunits by sucrose gradient fractionation, with the gray bars representing the portions of the peaks that were collected. **C.** SDS-PAGE analysis of wild-type 40S ribosomal subunits, 40S ribosomal subunits labeled with Cy5 via RACK1-ybbR (40S-ybbR-Cy5), as well as recombinant wild-type RACK1 and RACK1-ybbR. The bands highlighted in red boxes correspond to the proteins identified by mass mapping analysis in panel e. **D.** Western-blot analyses of the indicated purified 40S ribosomal subunits with select antibodies targeting proteins know to be involved in regulatory ribosomal protein ubiquitination. Asterix (***) indicate higher MW protein species, likely due to ribosomal protein modifications. The faint band below the RACK1-ybbR band may be due to partial cleavage of the ybbR tag, which is not labeled by SFP synthase. **E.** Native gel electrophoresis analysis of wild-type or 40S-RACK1-ybbR-Cy3 for binding to the HCV IRES (Cy5), and forming 40S-IRES and 80S-IRES complexes. Complexes were formed with the indicated ribosomal subunits at 2-fold excess to the HCV IRES, and the indicated Mg^2+^ concentration.

**Supplementary Figure 4 (related to Figure 3).**
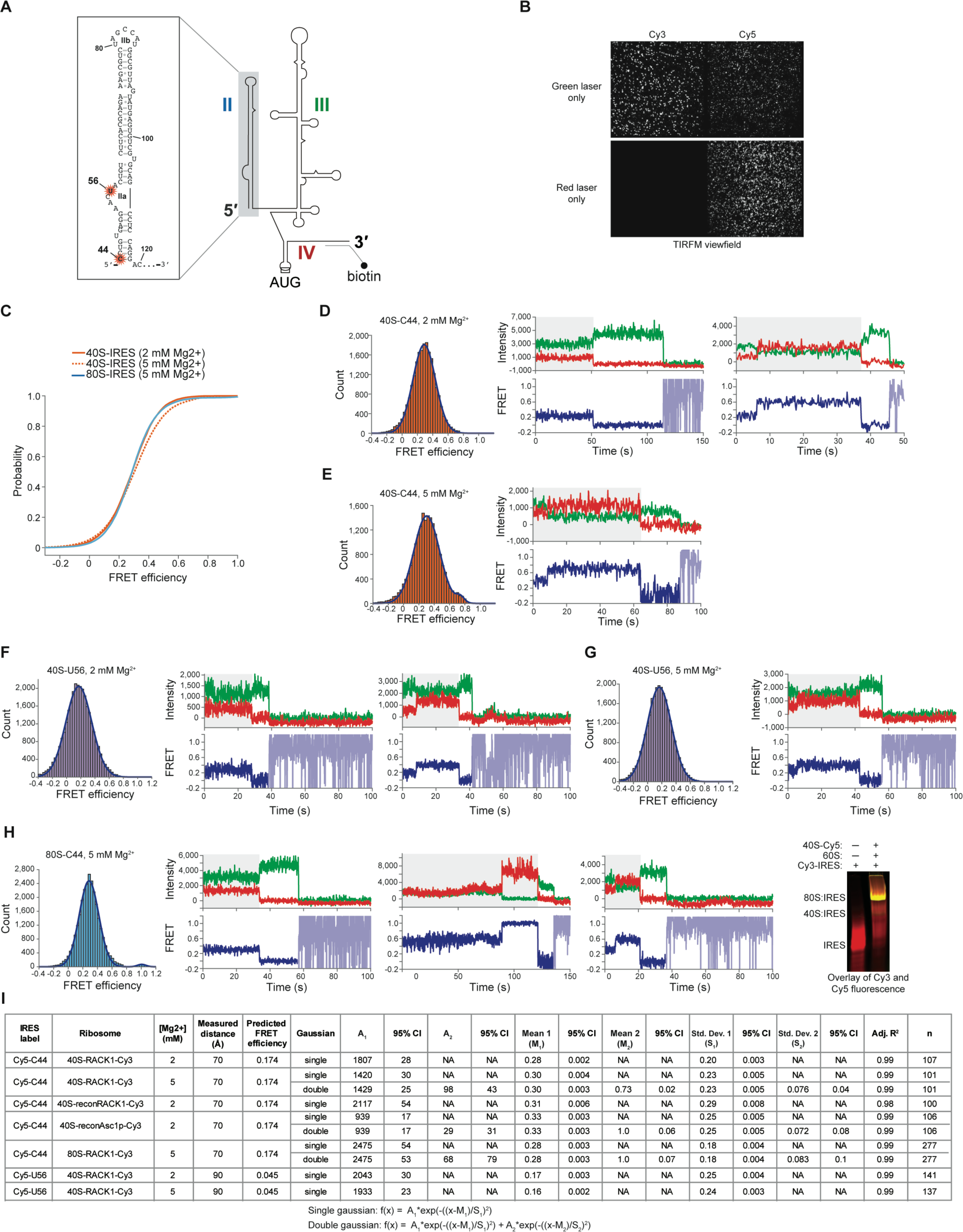
RACK1-ybbR is proximal to domain II of the HCV IRES and provides a signal for 40S-IRES conformational dynamics. **A.** Schematic of the secondary structure of the HCV IRES indicating domains II, III, and IV. Inset box shows an expansion of domain II with secondary structure of nucleotides, indicating positions of fluorescent labels at C44 and U56. An extension at the very 3’ end of the HCV IRES RNA was used to anneal a biotinylated oligo for immobilization in all single-molecule experiments. **B.** Example view field from TIRFM analyses of the 40S-RACK1-ybbR-Cy3:Cy5-IRES(C44) complex. **C.** Cumulative distribution plot of observed FRET intensities for RACK1-ybbR-Cy3 bound to IRES(Cy5-C44) (purple) either in a 40S-IRES complex in the presence of 2 mM Mg^2+^ (solid orange line) or 5 mM Mg^2+^ (orange dotted line), or in an 80S-IRES complex (solid blue line). **D.** FRET intensity histogram for 40S-RACK1-ybbR-Cy3 FRET with IRES(Cy3-C44) at 2 mM Mg^2+^ and representative single-molecule fluorescence traces. FRET intensity data were fit with a single Gaussian distribution. **E.** FRET intensity histogram for 40S-RACK1-ybbR-Cy3 FRET with IRES(Cy3-C44) at 5 mM Mg^2+^ and a representative single-molecule fluorescence trace. FRET intensity data was fit with a double Gaussian distribution. **F.** FRET intensity histogram for 40S-RACK1-ybbR-Cy3 FRET with IRES(Cy3-U56) at 2 mM Mg^2+^ and representative single-molecule fluorescence traces. FRET intensity data were fit with a single Gaussian distribution. **G.** FRET intensity histogram for 40S-RACK1-ybbR-Cy3 FRET with IRES(Cy3-U56) at 5 mM Mg^2+^ and representative single-molecule fluorescence traces. FRET intensity data were fit with a single Gaussian distribution. **H.** FRET intensity histogram for 80S-RACK1-ybbR-Cy3 FRET with IRES(Cy3-C44) at 5 mM Mg^2+^ and representative single-molecule fluorescence traces. FRET intensity data were fit with a double Gaussian distribution. After collecting TIRFM data, excess complex was analyzed by native gel electrophoresis to verify successful 80S formation (gel on right). Cy3 and Cy5 overlay used to indicate major co-localized fluorescent species. **I.** Table of parameters yielded from fitting the indicated experiments to single or double Gaussian distributions. Also showing distance measurements between labeling positions using PDB 5a2q and predicted FRET efficiencies for the Cy3-Cy5 dye pair.

**Supplementary Figure 5 (related to Figure 4).**
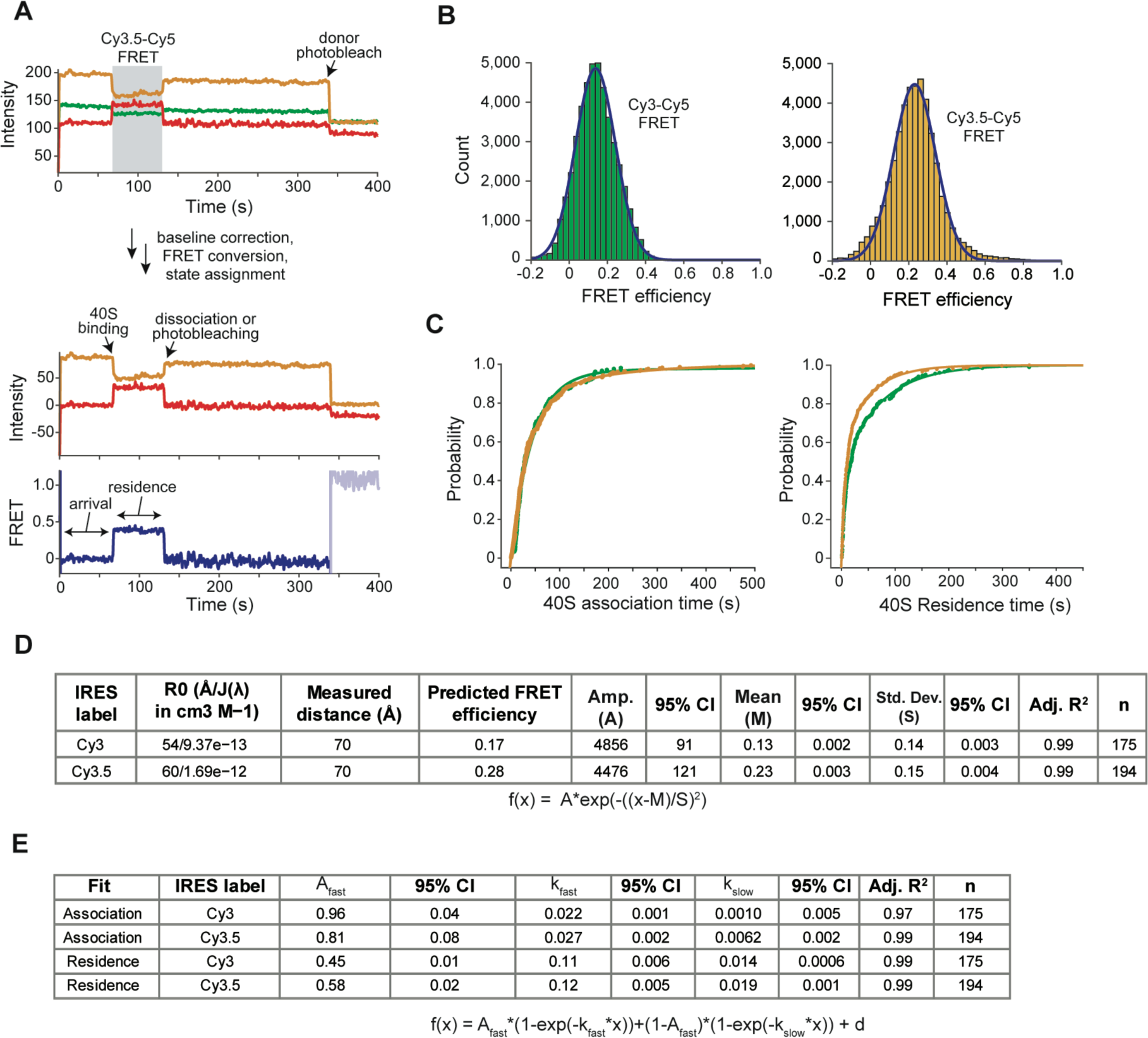
Monitoring 40S-IRES association kinetics by real-time smFRET in ZMWs. **A.** Example single-molecule fluorescence trace that depicts Cy3.5-Cy5 FRET upon delivery of 40S-RACK1-ybbR-Cy5 to dual-immobilized IRES(Cy3-C44) and IRES(Cy3.5-C44). The trace is displayed as in Figure 4b, and schematic used to show the process for extracting FRET efficiency and kinetic parameters. Briefly, ZMWs with either a single Cy3- or Cy3.5-IRES(C44) molecule were identified by a single-step photobleaching event of the donor fluorophore, based on their relative intensities. The respective FRET events are highlighted in grey, and photobleaching of the donors are indicated by the arrows. After selecting single molecules for each dye pair, we corrected for background fluorescence in each optical channel and calculated distributions of observed smFRET efficiencies for the Cy3-Cy5 and Cy3.5-Cy5 dye pairs. **B.** FRET intensity histograms for IRES(Cy3-C44, left green bars) or IRES(Cy3.5-C44, right orange bars) with 40S-RACK1-ybbR-Cy5. FRET intensity data were fit with single Gaussian distributions. Data is as in Figure 4c, but the histograms are now separated for clarity. **C.** Cumulative distribution plot of observed 40S association and residence times upon delivery to dual-immobilized IRESs. Colors reflect the separate Cy3 and Cy3.5 IRES data as in Figure 3. The lines represent fits to double-exponential functions with all fitting parameters in panel e. Given that the residence time of the 40S ribosomal subunit is concentration-independent, we attributed the minor difference in observed residence times with the two smFRET pairs to their distinct photophysical properties. **D.** Table of parameters yielded from fitting the indicated experiments to single or double Gaussian distributions. Also showing distance measurements between labeling positions using PDB 5a2q and predicted FRET efficiencies for the Cy3-Cy5 and Cy3.5-Cy5 dye pairs. **E.** Table of parameters yielded from fitting the indicated experiments to double-exponential functions.

**Supplementary Figure 6 (related to Figure 5).**
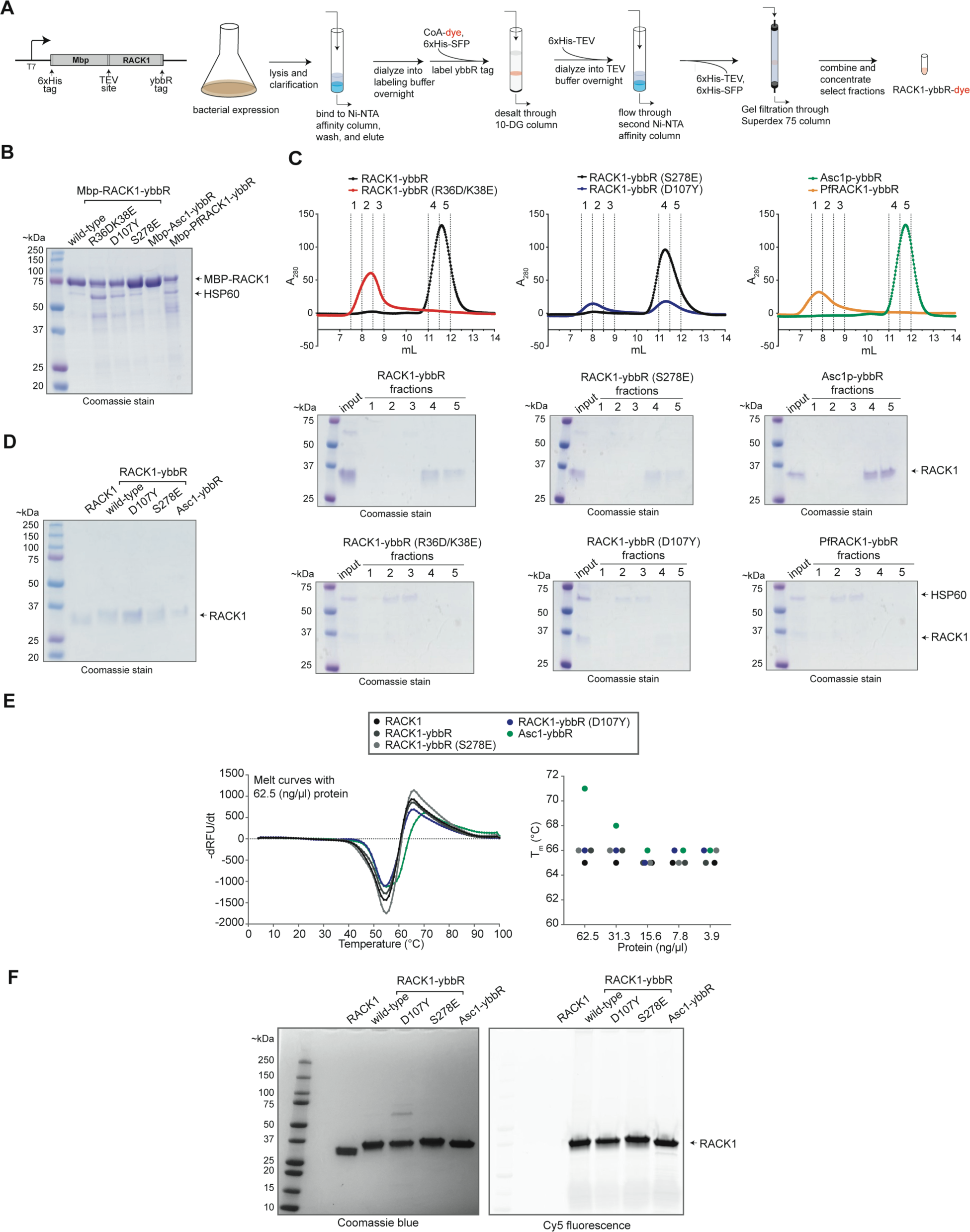
Purification and biochemical characterization of ybbR-tagged RACK1 mutants and homologs. **A.** Strategy to purify and fluorescently label recombinant RACK1-ybbR proteins. **B.** SDS-PAGE analysis of unlabeled 6xHis-Mbp-RACK1-ybbR proteins following first Ni-NTA purification and before TEV cleavage to remove MBP tag. **C.** Gel filtration profiles (Superdex 75) of unlabeled and ybbR-tagged RACK1 proteins following TEV cleavage and removal of tag by second Ni-NTA column. SDS-PAGE analysis of select fractions shows the presence of HSP60-bound RACK1 proteins in the void volume for select mutants (R36D/K38E and D107Y) and homologs (*Plasmodium falciparum* PfRACK1), while monomeric RACK1 elutes at ∼11 mL. **D.** SDS-PAGE of purified and unlabeled RACK1 proteins after concentration of fractions that elute at 11-12 mL for each protein. **E.** Thermal melt curves of various RACK1 proteins. The left panel shows the derivative of the relative fluorescence units (RFU) for melting of all proteins at 62.5 ng/μL. The right panel shows the estimated melting temperature (T_m_) for each protein at five concentrations. **F.** SDS-PAGE analysis of purified, concentrated, and Cy5-labeled RACK1-ybbR proteins. Both images are of the same gel, which was first scanned for Cy5 fluorescence (right) and subsequently stained with Coomassie blue (left). The band at ∼60kD in the D107Y mutant corresponds to HSP60.

**Supplementary Figure 7 (related to Figure 5).**
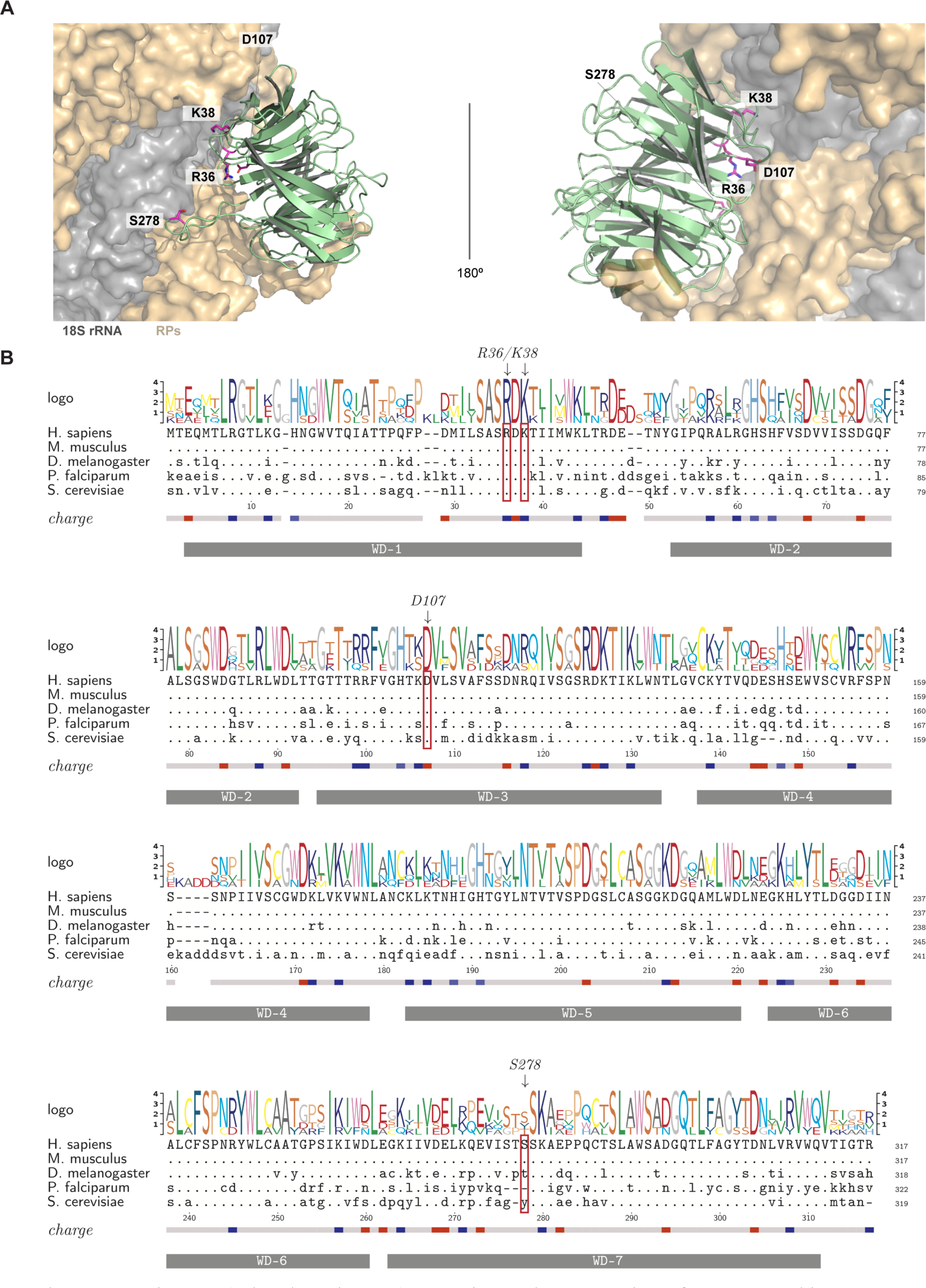
Location and conservation of RACK1 and its mutants. **A.** Model structure of RACK1 bound to the 40S ribosomal subunit (PDB: 5a2q), with the substituted residues indicated. The 18S rRNA is in grey, and ribosomal proteins are in tan. **B.** Alignment of the protein sequences for the indicated RACK1 orthologs. The residues boxed with red frames are those that are indicated in panel a.

**Supplementary Figure 8 (related to Figure 5).**
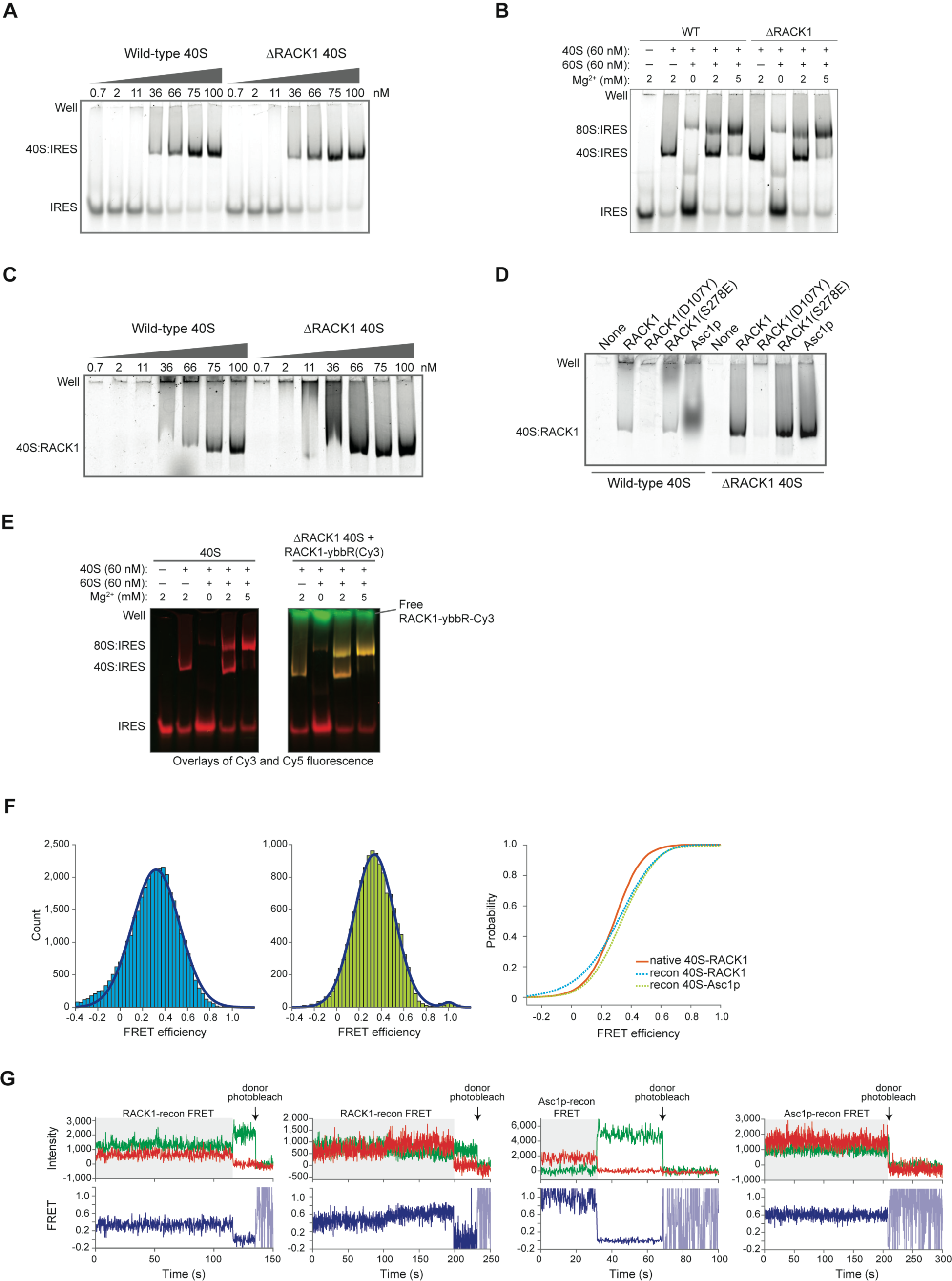
*In vitro* reconstitution of RACK1 to study RACK1 association with human ribosomes. **A.** Native gel analysis of the indicated 40S ribosomal subunits binding to Cy5-labeled HCV IRES. **B.** Native gel analysis of wild-type and ΔRACK1 ribosomes forming 40S- and 80S-IRES complex on a Cy5-labeled HCV IRES. **C.** Native gel analysis of recombinant RACK1-ybbR-Cy5 incorporation into the indicated 40S ribosomal subunits. The concentration of RACK1-ybbR-Cy5 was 40 nM, and the concentration of 40S subunits are indicated above each lane. A representative gel is shown. **D.** Native gel analysis of recombinant RACK1-ybbR-Cy5, RACK1(D107Y)-ybbR-Cy5, RACK1(S278E)-ybbR-Cy5, and Asc1p-ybbR-Cy5 incorporation into the indicated 40S ribosomal subunits (75 nM). The concentration of each protein was 40 nM. **E.** Native gel analysis of reconstituted 80S ribosomes binding to Cy5-labeled HCV IRES. Wild-type 40S and in vitro reconstituted ΔRACK1 40S:RACK1-ybbR-Cy3 were analyzed in parallel. The gel displays the overlay of Cy3 (green) and Cy5 (red) fluorescence signals. **F.** Cumulative distribution plot and histograms for FRET efficiency for reconstituted RACK1-ybbR-Cy3 (“RACK1 recon”) and Asc1p-ybbR-Cy3 (“Asc1p recon”) ribosomes versus the natively-purified 40S-RACK1-ybbR ribosomes. FRET intensity histograms show counts for the RACK1 and Asc1p reconstituted ribosomes on the left (blue) and right (green), respectively. The number of traces analyzed are indicated in Supplemental Figure 4i. **G.** Example single-molecule fluorescence traces that captured FRET between the indicated 40S ribosomal subunits labeled with Cy3 and HCV IRES(C44-Cy5). The FRET events are highlighted in gray, and photobleaching of the Cy3 donor dye is indicated by the arrow.

**Supplementary Figure 9 (related to Figure 5).**
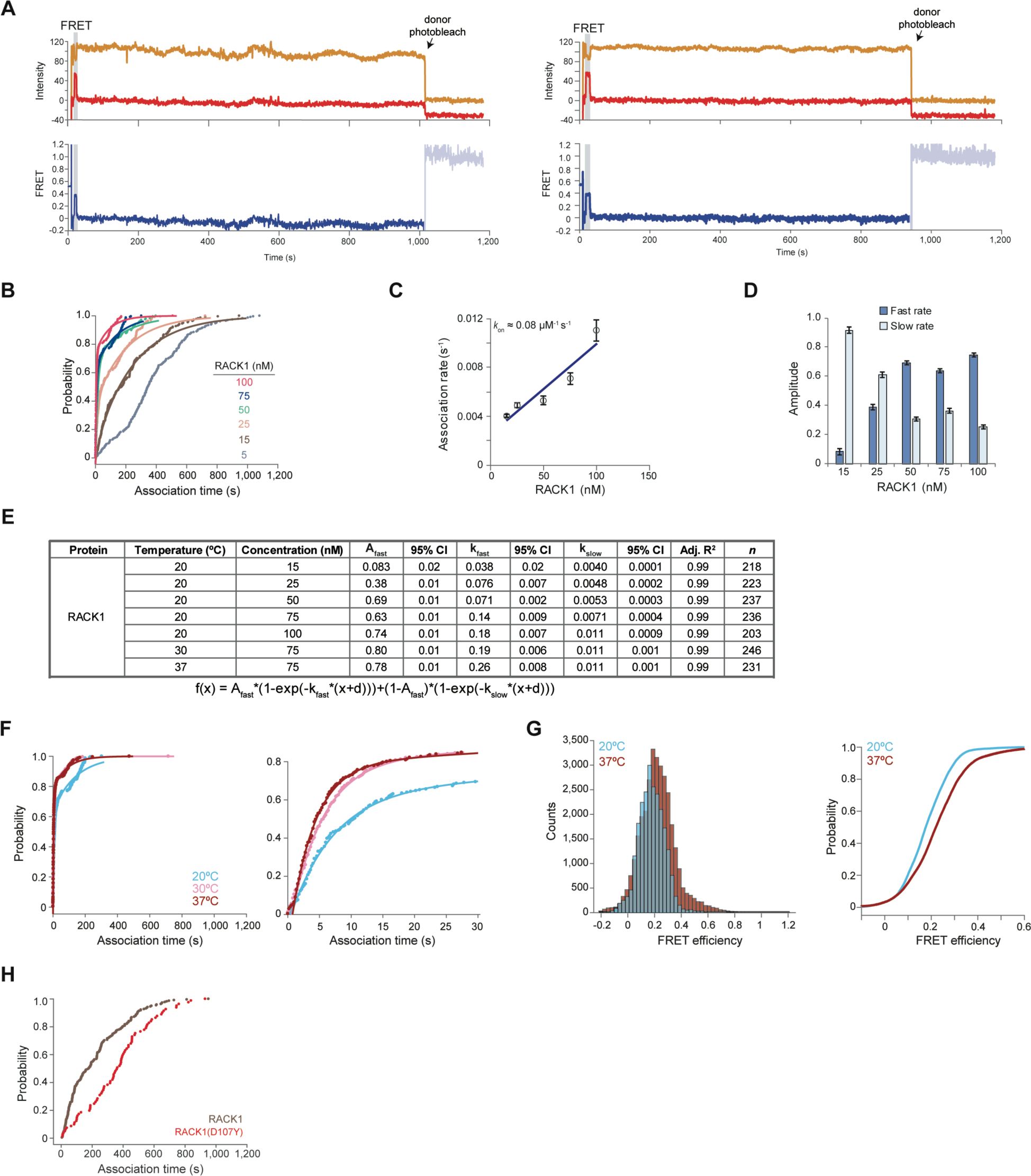
Real-time monitoring of RACK1 association to single ribosomes by smFRET in ZMWs. **A.** Example single-molecule fluorescence traces that captured FRET between RACK1 labeled with Cy5 and the immobilized 40S-HCV IRES(C44-Cy3.5) complex. The FRET events are highlighted in gray, and photobleaching of the Cy3.5 donor dye is indicated by the arrow. **B.** Cumulative distribution plot of observed RACK1 association times from 0 to 1,200 seconds. The labels indicate RACK1-ybbR-Cy5 concentration (nM), and the lines represent fits to double-exponential functions. **C.** Plot of observed RACK1 association rates (*k_obs_*) of the slow phase at the indicated concentrations. Error bars represent the 95% confidence interval for each value. The line represents a fit to a linear function, with equation y = 0.00008x + 0.0025 and *R^2^* = 0.8889, which yielded the indicated association rate (*k_on_*) for the slow phase. **D.** Plot of the calculated amplitudes for the fast and slow phases yielded from fits of the indicated datasets to double-exponential functions. **E.** Table of parameters yielded from fitting the indicated experiments to double-exponential functions. **F.** Cumulative distribution plots of observed RACK1 association times from 0 to 1,200 (left) and 0 to 30 (right) seconds. RACK1-ybbR-Cy5 was delivered at 75 nM at the indicated temperatures. The lines represent fits to double-exponential functions. **G.** Histogram (left) and cumulative distribution plots of observed FRET efficiencies between the HCV IRES-Cy3.5 (donor) and RACK1-ybbR-Cy5 (acceptor) upon RACK1 binding to the immobilized ΔRACK1 40S-IRES complex at the indicated temperatures. **H.** Cumulative distribution plot of observed RACK1(D107Y) association times from 0 to 1,200 seconds after delivery at 15 nM. We replotted 15 nM RACK1-ybbR-Cy5 data from panel a here as a comparison.

**Supplementary Figure 10 (related to Figure 5).**
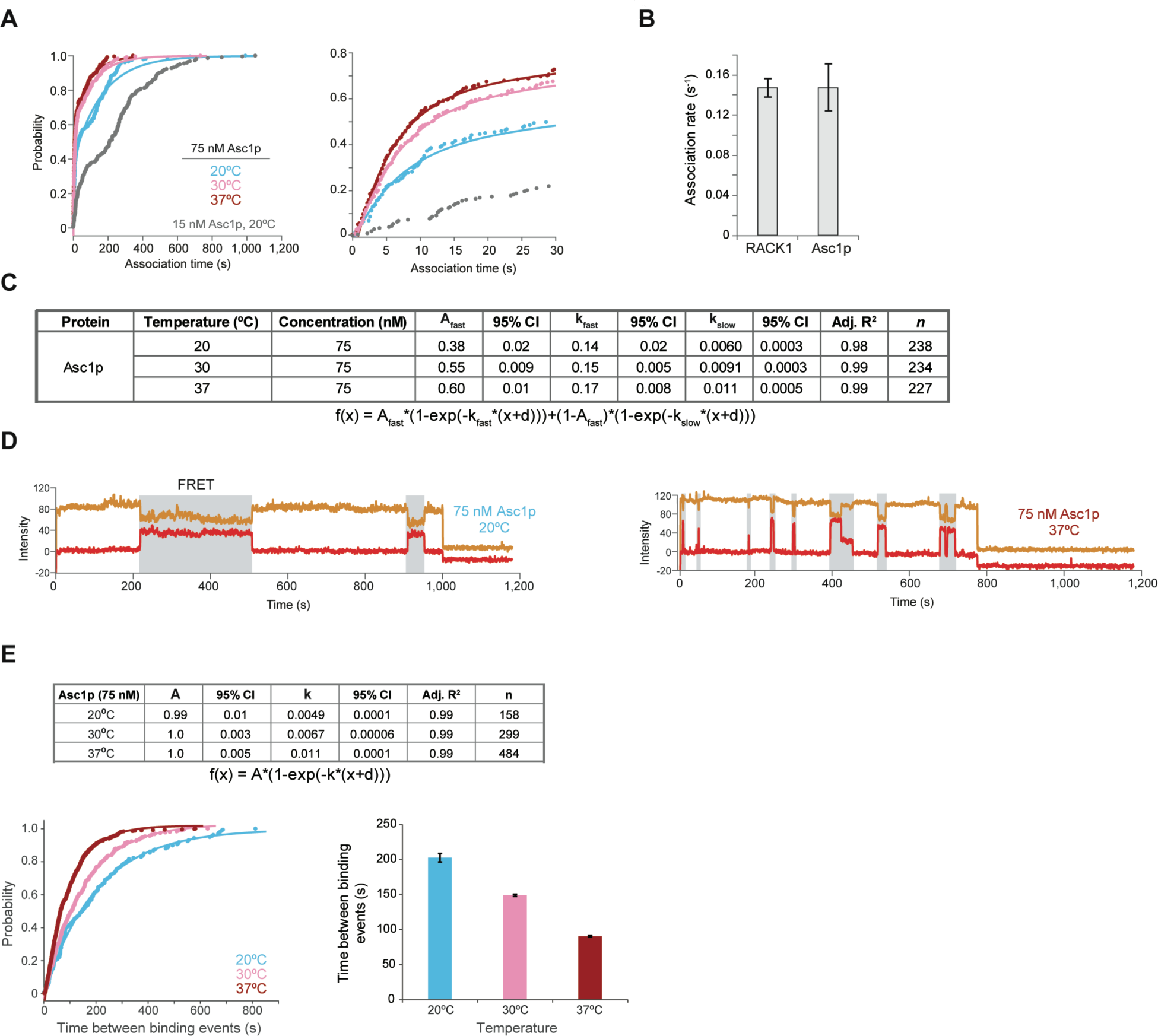
Dynamic association of *S. cerevisiae* Asc1p with 40S human ribosomal subunits measured by real-time smFRET in ZMWs. **A.** Cumulative distribution plots of observed Asc1p association times from 0 to 1,200 (left) and 0 to 30 (right) seconds. Asc1p-ybbR-Cy5 was delivered at 75 nM at the indicated temperatures, as well as at 15 nM at 20°C. The lines represent fits to double-exponential functions. **B.** Plot of observed association rates (*k_obs_*) for the fast phase upon delivery of 75 nM Asc1p-ybbR-Cy5 or RACK1-ybbR-Cy5 at 20°C. Error bars represent the 95% confidence interval for each value. **C.** Table of parameters yielded from fitting the indicated experiments to double-exponential functions. **D.** Representative single-molecule fluorescence traces for Asc1p-ybbR-Cy5 binding to a single ΔRACK1 40S:IRES-Cy3.5 complex indicated by Cy3.5-Cy5 FRET, highlighted in gray. Single 40S-IRES complexes were identified via a single-step photobleaching event in the Cy3.5 donor signal, indicated by the arrow. **E.** The time between two Asc1p binding events on a single 40S-IRES complex was measured upon its delivery at 75 nM at the indicated temperatures. The table (top) contains the parameters from fits to single-exponential functions, with *n* representing the number of binding events that were analyzed. The cumulative distribution plot (bottom left) displays the observed time between Asc1p binding events on a single 40S-IRES complex and the lines represent the fits to single-exponential functions. The bar plot (bottom right) displays the calculated times between Asc1p binding events for the indicated experiments yielded from the fits to single-exponential functions, with error bars representing the 95% confidence interval for each value.

**Supplementary Figure 11 (related to Figure 5).**
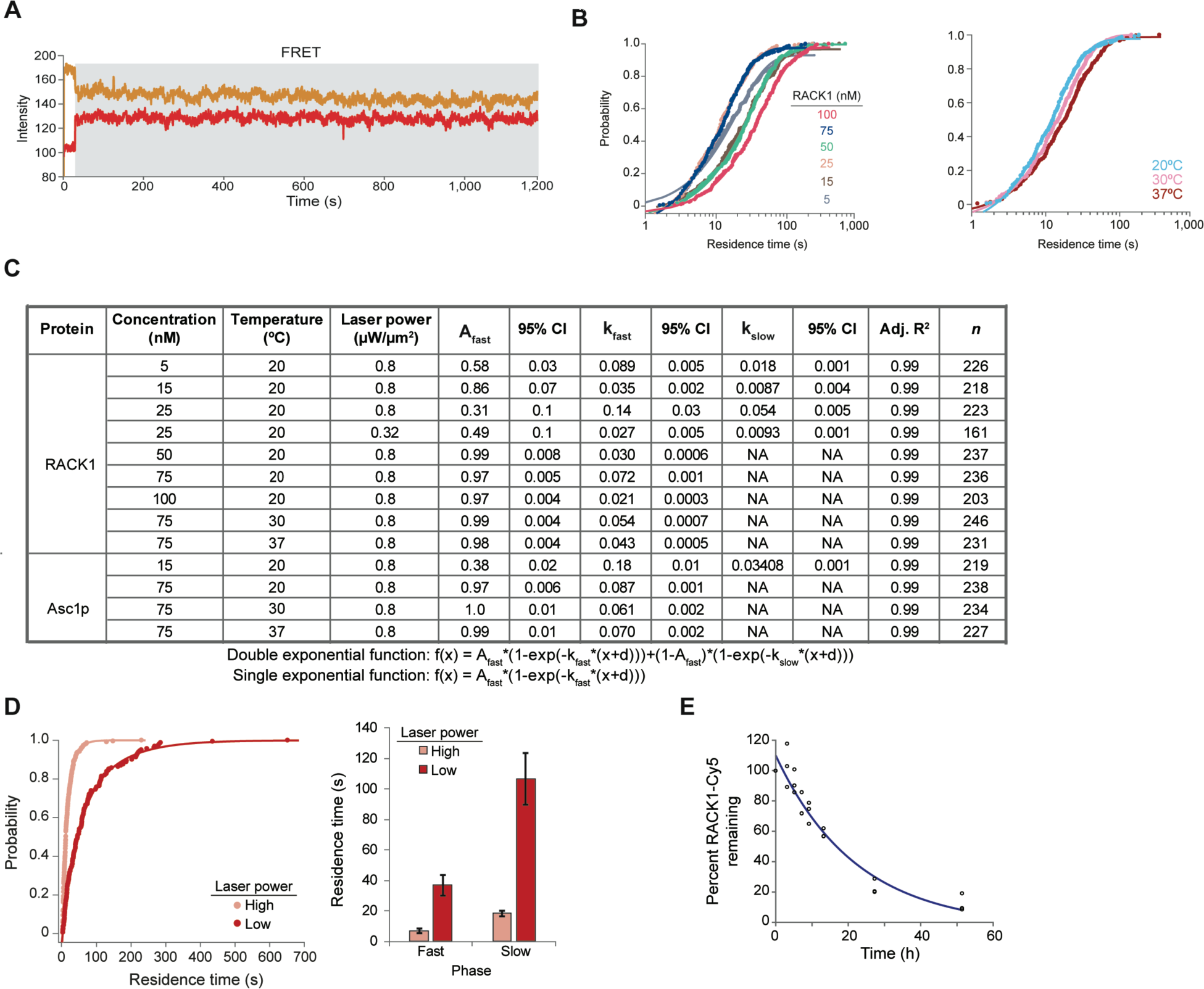
RACK1 dissociation measurement by smFRET in ZMWs is limited by photobleaching of the dyes. **A.** A single-molecule fluorescence trace (prior to background correction) for RACK1-ybbR-Cy5 binding to a single ΔRACK1 40S:IRES-Cy3.5 complex indicated by Cy3.5-Cy5 FRET, highlighted in grey. While rare due to the limited lifetime of the Cy5 dye, we observed RACK1 binding events that lasted greater than 10 minutes. **B.** Cumulative distribution plots of observed RACK1 residence times at the indicated concentrations (left) and temperatures (right, delivery at 75 nM). The lines represent fits to single- or double-exponential functions, as indicated in panel c. **C.** Table of parameters yielded from fitting the indicated experiments to single- or double-exponential functions. “NA” indicates the respective parameters that were not calculated since these experiments were best fit by single-exponential functions. The respective equation is listed below the table. **D.** Cumulative distribution (left) and column (right) plot of observed residence times for RACK1 after delivery at 15 nM using low (0.32 µW/ µm^2^) or high (0.8 µW/ µm^2^) laser power. Error bars represent the 95% confidence interval for each value. **E.** Native gel electrophoresis analysis of 40S-RACK1-ybbR-Cy5 following competition after the indicated times with recombinant unlabeled RACK1-ybbR at 20-fold excess. Each circle on the graph represents the fluorescence intensity remaining relative to the 0.08 h time point from each replicate, which were used to generate the plot in Figure 5h. The line represents the fit to an exponential function.

